# Tutorial on MUedit: An open-source software for identifying and analysing the discharge timing of motor units from electromyographic signals

**DOI:** 10.1101/2023.07.13.548568

**Authors:** Simon Avrillon, François Hug, Ciara Gibbs, Dario Farina

**Affiliations:** Department of Bioengineering, Faculty of Engineering, Imperial College London, London W12 7TA, UK; Université Côte d’Azur, LAMHESS, Nice 06200, France; The University of Queensland, School of Biomedical Sciences, St Lucia QLD 4072, QLD, Australia

## Abstract

We introduce the open-source software MUedit and we describe its use for identifying the discharge timing of motor units from all types of electromyographic (EMG) signals recorded with multi-channel systems. MUedit performs EMG decomposition using a blind-source separation approach. Following this, users can display the estimated motor unit pulse trains and inspect the accuracy of the automatic detection of discharge times. When necessary, users can correct the automatic detection of discharge times and recalculate the motor unit pulse train with an updated separation vector. Here, we provide an open-source software and a tutorial that guides the user through i) the parameters and steps of the decomposition algorithm, and ii) the manual editing of motor unit pulse trains. Further, we provide simulated and experimental EMG signals recorded with grids of surface electrodes and intramuscular electrode arrays to benchmark the performance of MUedit. Finally, we discuss advantages and limitations of the blind-source separation approach for the study of motor unit behaviour during muscle contractions in humans.

## Introduction

The nervous system transforms a motor intent into movements by transmitting electrical action potentials to contractile tissues, which ultimately results in force production. The functional unit in charge of the conversion of this electrical activity into a mechanical output is the motor unit, which comprises an alpha motor neuron and the muscle fibres it innervates (Liddell and Sherrington, 1925). Soon after the description of the motor unit, experimentalists developed tools to record their extracellular electrical activity, with aim of understanding how the nervous system generates and controls muscle force. From concentric needles (Adrian and Bronk, 1929) to fine wires (Basmajian and Stecko, 1962), from large surface electrodes (de Luca, 1997, Farina et al., 2004) to dense grids of small surface electrodes (Farina et al., 2016), the set of tools available to assess motor unit activity has expanded over the years (Duchateau and Enoka, 2011). The recent advent of grids of surface electrodes (Farina et al., 2016) and intramuscular electrode arrays (Chung et al., 2023, Farina et al., 2008b, Muceli et al., 2022, Muceli et al., 2015), together with algorithms that automatically separate the overlapping activity of motor units [for recent examples, see (Chen and Zhou, 2016, Holobar and Zazula, 2007, McGill et al., 2005, Nawab et al., 2008, Negro et al., 2016)], has ushered the study of the neural control of movement into a new era.

While progress of EMG decomposition over the last decades has been remarkable, many of the approaches have not yet been standardised. For example, the number of identified motor units and the accuracy of the detection of their discharge timing strongly depend on the decomposition algorithm used, but there is no consensus on the optimal algorithm. The situation is exacerbated by the absence of open-source software; most decomposition algorithms used by the community are compiled in proprietary hardware and software (e.g., OT BioLab, Delsys, Demuse). Therefore, the constituents of these algorithms cannot be inspected or tested by their users. Other decomposition algorithms have been implemented and shared by research groups, but without user interfaces that allow experimentalists to easily replicate their results (Formento et al., 2021, Jiang et al., 2021). Importantly, each of these decomposition algorithms should be associated to an interactive tool that displays the result of the automatic decomposition and enables users to correct it when performing neurophysiological investigations. Recent tutorials have emphasised the importance of manual editing with guidance to standardise it (Del Vecchio et al., 2020, Hug et al., 2021), but without an open-source software that permits global application.

We introduce an open-source software for identifying and analysing the discharge timing of motor units from electromyographic (EMG) signals. We also provide here a tutorial and data samples to carefully guide researchers through the steps of EMG decomposition. After validating the algorithm with simulated surface and intramuscular EMG signals, we present the different steps of the algorithm with various options that can impact the results. The software also offers capabilities to inspect and manually correct the results of the decomposition. We report in this article the main steps of manual editing to guarantee an accurate detection of the motor unit discharge times. All the material (software, code, user manual, signals) is available at https://github.com/simonavrillon/MUedit.

## Decomposition of EMG signals

We will first summarise the main principles of the decomposition of EMG signals. Intramuscular electrodes have a high spatial selectivity, such that each recording comprises the activity of one to a few motor units. In this case, a human operator can visually identify and sort the unique profiles of the action potentials of the detected units. Automatic algorithms can perform the same process without the need for lengthy manual analyses and, in some cases, can also separate action potentials that overlap in time (LeFever and De Luca, 1982, McGill et al., 2005, Nawab et al., 2008). Most algorithms for intramuscular EMG decomposition are based on detection and classification of action potential waveforms, a process called spike sorting.

Surface electrodes are less selective than intramuscular electrodes. While their spatial selectivity can be increased by high-pass spatial filtering, such as the Laplacian filter, conventional spike sorting methods developed for intramuscular EMG do not usually achieve sufficiently high accuracy when decomposing surface EMG (Gazzoni et al., 2004, Rau et al., 1997). On the other hand, EMG decomposition can be also achieved by inverting the signal generation model, an operation that is particularly successful when EMG signals are recorded with a large number of electrodes (Figure 1) (Farina and Holobar, 2016, Holobar and Farina, 2014). The inversion of the EMG generation model is equivalent to the application of a source separation algorithm, such as the fast Independent Component Analysis [fastICA; (Chen and Zhou, 2016, Negro et al., 2016)] or the Convolution Kernel Compensation approach [CKC; (Holobar and Zazula, 2007)]. In short, these methods iteratively optimise separation vectors for each motor unit to separate its series of discharge times from all other active motor units (Farina and Holobar, 2016, Holobar and Farina, 2014). The projection of the pre-processed EMG signals onto these separation vectors estimates motor unit pulse trains, from which one can detect the discharge times (Figure 2). Interestingly, the generative model of EMG signals is identical for surface and intramuscular signals, though the bandwidth of action potential waveforms differs. Therefore, source separation algorithms can be applied to both types of signals, as illustrated in Figure 2.

**Figure 1.**
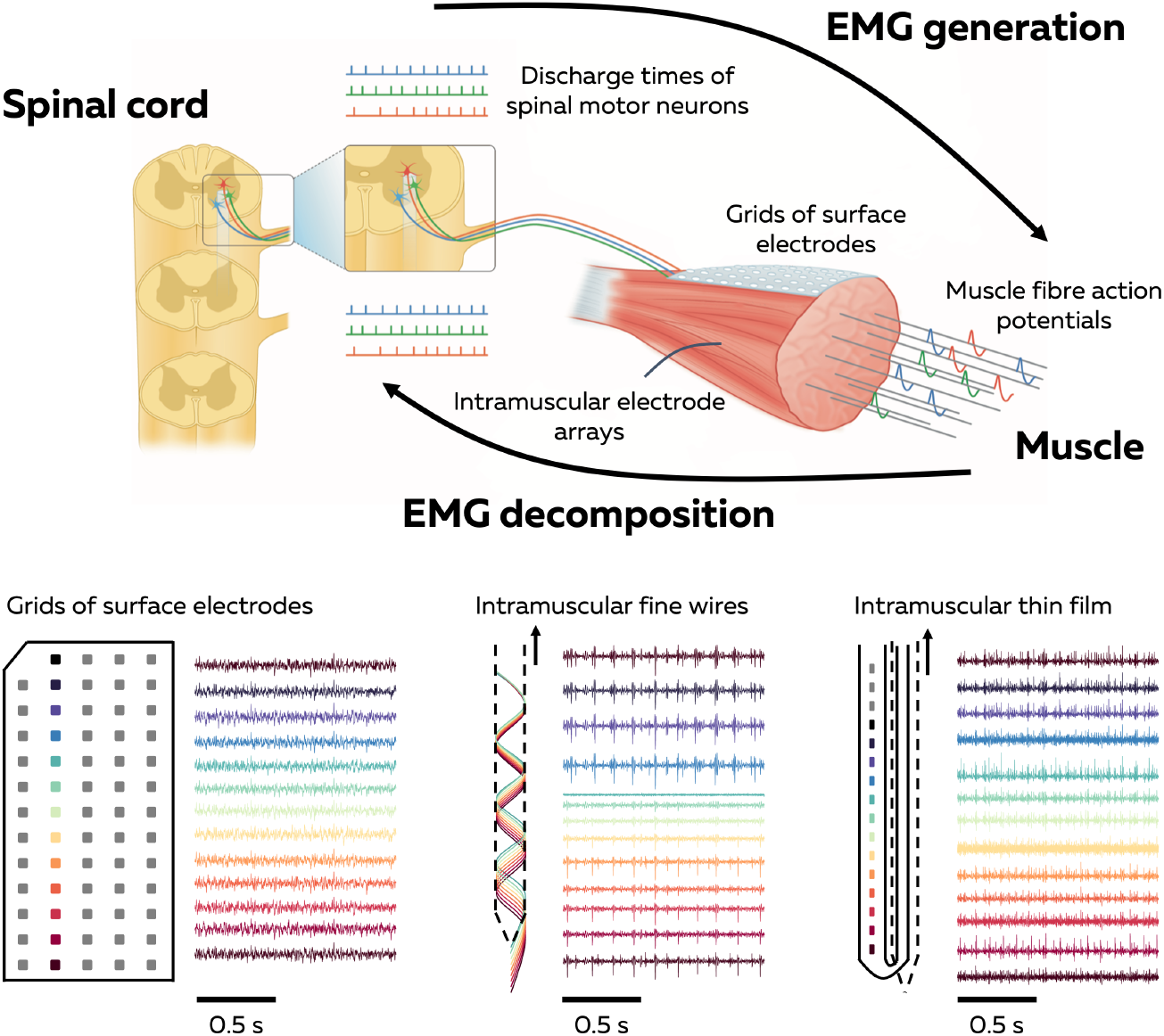
Overview of the EMG decomposition. The nervous system generates force by activating motor units and modulating their discharge rates. During isometric contractions, the train of motor unit action potentials can be modelled as the convolution of the series of discharge times (mathematically represented as Dirac delta functions) and the motor unit action potentials. The EMG signals are the sum of the trains of action potentials from the active motor units within the recorded muscle volume. The EMG decomposition with blind source separation consists of inverting the generative model of EMG signals by estimating the series of discharge times for active motor units. For this, blind-source separation algorithms can be applied on EMG signals recorded with grids of surface electrodes or intramuscular electrode arrays. Representative samples of these EMG signals are displayed for 13 adjacent electrodes.

**Figure 2.**
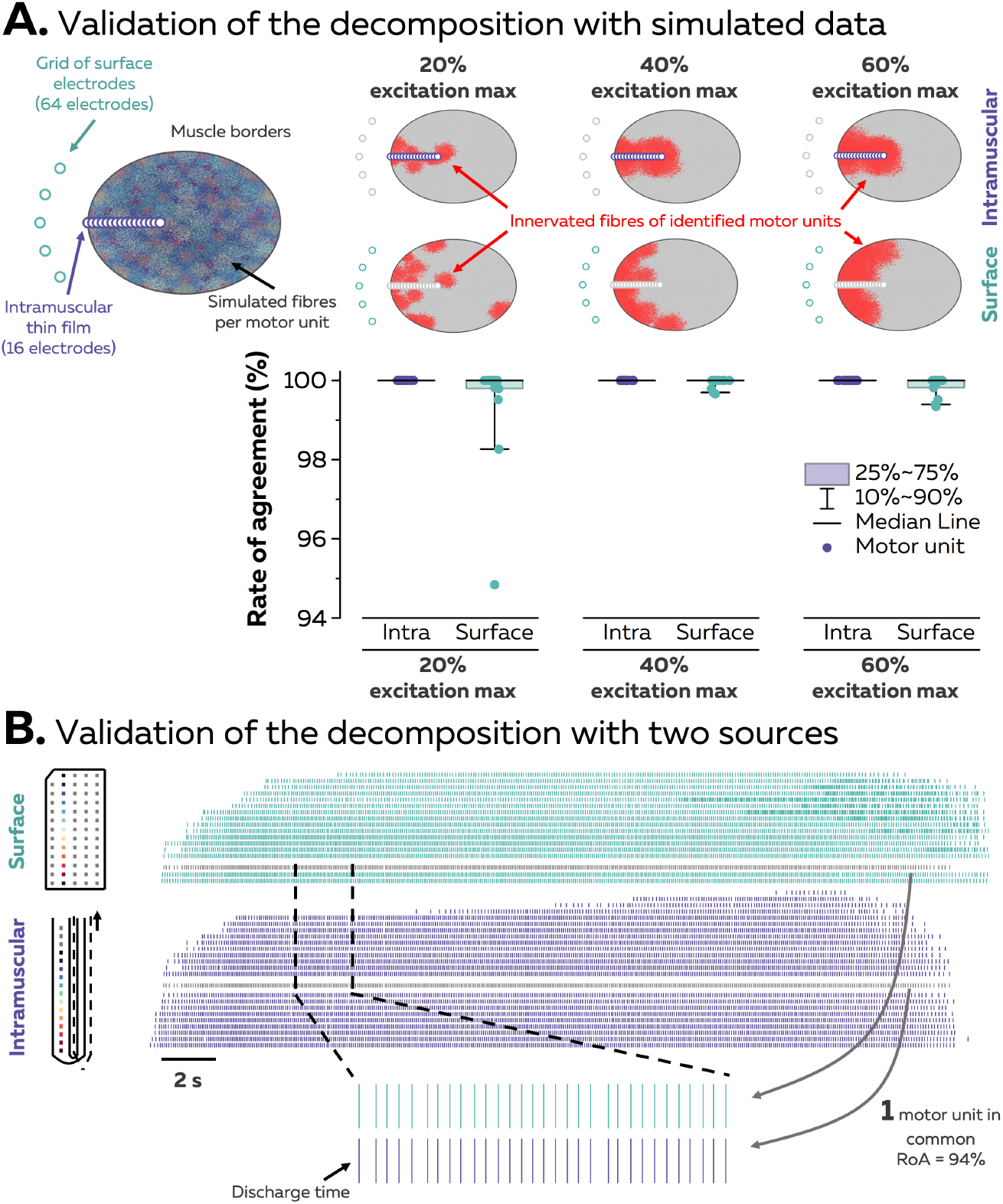
Validation of the EMG decomposition. We validated the decomposition algorithm implemented in the MUedit software using simulations of 200 motor units active for relative levels of 20%, 40%, and 60% of the maximal excitation. The territories of their innervated fibres are displayed with different colours on a transverse view of the muscle (A). The simulated EMG signals were either detected by a grid of 64 surface electrodes or an array of 16 intramuscular electrodes (A). Then, we calculated the rate of agreement between the series of discharge times identified by the decomposition algorithm and the ground truth (the simulated series of discharge times). Above the graph, the position of the innervated fibres from identified motor units are displayed in red. We also compared the motor units identified from both surface and intramuscular (thin film) EMG signals concurrently recorded from the Tibialis Anterior muscle during an experiment (B). We only identified one motor unit in common between both recordings, with a rate of agreement of 94% after edition. Each tick is a discharge time.

Our software – MUedit – uses the fast independent component analysis (ICA) to estimate the pulse trains and detect the discharge times of individual motor units. This method has already been theoretically described (Farina and Holobar, 2016) and validated (Negro et al., 2016). Here, we provide a documented implementation of the open-source algorithm with various options to standardise the usage of EMG decomposition. For this, we first simulated EMG signals (surface and intramuscular) that prospective users can download to verify that they correctly operate the software (Figure 2). Following this, we estimate the rate of agreement between simulated and estimated series of motor unit discharge times, calculated as the ratio between correctly identified discharge times and the sum of correctly identified discharge times, missed discharge times, and falsely identified discharge times.

For the simulated EMG signals, we generated the series of discharge times of a group of 200 motor units in an anatomical model entailing a cylindrical muscle volume with parallel fibres (Farina et al., 2008a, Konstantin et al., 2020), in which subcutaneous and skin layers separate the muscle from the surface electrodes (Figure 2). We set the radius of the muscle to 25.4 mm and the thicknesses of the subcutaneous and skin layers to 5 mm and 1 mm, respectively. The centres of the motor units were distributed within the cross section of the muscle using a farthest point sampling technique. The farthest point sampling filled the cross-section by iteratively adding centre points that were maximally distant from all the previously generated motor unit centres, resulting in a random and even distribution of the motor unit territories within the muscle (Figure 2). The number of fibres innervated by each motor neuron followed an exponential distribution, ranging from 15 to 1500. The fibres of the same motor unit were positioned around the centre of the motor unit within a radius of 0.5 to 27.7 mm, and a density of 20 fibres/mm^2^ (Armstrong et al., 1988). Because motor unit territories were intermingled, the density of fibres in the muscle reached 200 fibres/mm^2^. The motor unit action potentials were detected in the model by either a grid of 64 circular surface electrodes with a diameter of 1 mm arranged in 5 columns and 13 rows (inter-electrode distance: 4mm), or 16 circular intramuscular electrodes arranged in a single array (inter-electrode distance: 1mm). The grid was centred over the muscle in the transverse direction. The intramuscular electrode array was centred respective to the grid and inserted perpendicular to the longitudinal direction of the volume toward the centre of the muscle (Figure 2). After automatic decomposition and manual edition, the rates of agreement were above 99% (median 100%) and 95% (median 100%) for intramuscular and surface EMG signals during contractions at relative intensities of 20%, 40%, and 60% of the maximal excitation (Figure 2). These values are comparable to those obtained in previous validation studies of the fastICA algorithm (Negro et al., 2016). MUedit thus provides an accurate estimation of series of motor unit discharge times. The decomposition of these simulated signals and the parameters of the decomposition are available at https://github.com/simonavrillon/MUedit.

## Recording and pre-processing of EMG signals

The application of blind source separation algorithms on EMG signals relies on specific theoretical and practical considerations. First, the number of observations (electrodes) should always be greater than the number of identifiable motor units. Although it is likely that the number of active motor units is often higher than the number of electrodes, we consider as identifiable motor units those with action potentials accounting for most of the energy of the EMG signals (Farina and Holobar, 2016, Holobar and Farina, 2014). These motor units are either close to the electrodes or are composed by many muscle fibres within the recorded muscle volume. All other motor units contribute to the noise component of the model. Second, identifiable motor units must have spatio-temporal action potential waveforms that differ from all other active motor units within the recorded volume. In practice, this means that motor units innervating a similar number of muscle fibres and positioned close to each other within the recorded muscle volume will be likely not distinguishable. Again, increasing the number and spatial distribution of recording electrodes increases the likelihood that each motor unit will have a unique motor unit action potential profile (Caillet et al., 2023, Farina et al., 2008a).

For this tutorial article, we used three EMG recording modalities (Figure 1). First, we recorded surface EMG signals from the tibialis anterior with two-dimensional adhesive grids of 64 electrodes [13 × 5 electrodes with one electrode absent on a corner, gold-coated, interelectrode distance 4 mm; (GR04MM1305, OT Bioelettronica, Italy)] positioned next to each other to create a grid of 256 electrodes separated by 4 mm. Before the placement of the grids, the skin was shaved and cleansed with an abrasive gel (Nuprep, Weaver and company, USA). Each adhesive grid was held on the skin using a semidisposable biadhesive foam layer. The cavities within the adhesive layer were filled with conductive paste (SpesMedica, Italy) to facilitate the skin-electrode contact. Second, we recorded intramuscular EMG signals from the tibialis anterior with an array of 16 fine wires (stainless steel wires coated with Teflon, insulated diameter: 50 μm, California wires, USA). Eight fine wires were threaded within a 23G hypodermic needle and bent at the end to form a hook. Two needles were inserted within the muscle tissue with approximate angles of 30 degrees and 45 degrees, respectively. The insertion was guided with a portable ultrasound probe (Butterfly IQ+, Butterfly Network, USA) to reach tissues above and below the central aponeurosis of the Tibialis Anterior muscle. Finally, we recorded intramuscular EMG signals from the tibialis anterior with an intramuscular linear array of 40 electrodes on a thin film [platinum coated, interelectrode distance: 0.5 mm; (Muceli et al., 2022)]. The thin film was inserted within the muscle tissue with a 25G hypodermic needle that pulled the thin film with a guiding filament. The insertion was guided with the same portable ultrasound probe to reach tissues above the central aponeurosis of the Tibialis Anterior. Two strap electrodes (ground and reference electrodes) dampened with water were placed around the ankle for each data collection. The EMG signals were all recorded in monopolar mode at a frequency of 2048 Hz for surface EMG signals and 10240 Hz for intramuscular EMG signals (EMG-Quattrocento, 400 channel EMG amplifier; OT Bioelettronica, Italy).

As depicted in Figure 2, the use of intramuscular and surface electrodes leads to the identification of different groups of motor units. Grids of surface electrodes cover a larger surface of the muscle, but mostly identify motor units positioned in the superficial portion of the muscle volume (Figure 2). Conversely, intramuscular electrode arrays identify motor units close to the array. Thus, during simultaneous recordings of surface and intramuscular EMG signals, we only identified a single common motor unit from separated decompositions of intramuscular (14 motor units from a thin film) and surface (17 motor units from a grid of 64 electrodes placed over the thin film) EMG signals (Figure 2).

The reliable identification of the entire series of discharge times for each motor unit requires stationary recording conditions to keep the separation vector consistent. In practice, this means that EMG signals should be recorded during isometric contractions as a change in position/orientation of the active muscle fibres relative to the electrodes during the recordings can impact temporal and spatial action potential profiles. While intramuscular electrode arrays may be less impacted by changes in muscle geometry, displacement of the electrodes relatively to the source can still occur due to electrode drifts within the tissue during dynamic contractions. For this tutorial article, data was collected during submaximal isometric dorsiflexions. Four participants (1 male and 1 female for the grids of surface electrodes, 1 male for the array of fine wires, 1 male for the thin film) sat on a chair with the hips flexed at 85°, 0° being the hip neutral position, and their knees fully extended. We fixed the foot of the dominant leg (right in all participants) onto the pedal of a commercial dynamometer (OT Bioelettronica, Turin, Italy) positioned at 30° in the plantarflexion direction, 0° being the foot perpendicular to the shank. The foot was fixed to the pedal with inextensible straps positioned around the proximal phalanx, metatarsal and cuneiform. Force signals were recorded with a load cell (CCT Transducer s.a.s, Turin, Italy) connected in-series to the pedal using the same system as for the EMG recordings. The dynamometer was positioned accordingly to the participant’s lower limb length and secured to a table to avoid any motion during the contractions.

EMG signals were imported into the MUedit software for decomposition. We visually inspected all the signals to remove the channels with artifacts or low signal to noise ratio. This is because the presence of artifacts and noise may promote convergence of the decomposition algorithm toward unreliable separation vectors. Signals were then band pass filtered using a 2^nd^ order Butterworth filter (20-500 Hz for surface EMG signals, 100-4400 Hz for intramuscular EMG signals). The filtered EMG signals were extended and whitened before the identification of separation vectors. The extension consisted of duplicating each EMG signal with a delay of 1 to *n-1* data samples in order to have a high ratio between the number of observation and the number of identifiable motor units, *n* being the extension factor. For example, with 64 EMG signals and an extension factor of 16, we obtained 1024 signals with 16 duplicates per signal shifted by 0 to 15 data samples. Negro et al. (2016) previously suggested that choosing an extension factor *n* to reach 1000 signals maximise the number of identified motor units. Following this, the extended EMG signals were demeaned and whitened to make them spatially independent and of equal power.

## Optimisation of motor unit separation vectors

After pre-processing of the EMG signals, the steps to follow consist of i) iteratively optimising separation vectors for each motor unit to identify a matrix that transform the whitened EMG signals into the motor unit pulse trains and ii) identifying the discharge times for each motor unit from these pulse trains (Figure 3A). Separation vectors can be initialised using the values of the EMG signals at the time instants where the value of the sum of squared EMG signals is maximised (Figure 3A). This ‘activity index’ aims at speeding up the convergence of the optimisation loop toward a reliable separation vector. Alternatively, the MUedit software introduces a second option where separation vectors are initialised with random weights following a Gaussian distribution (Figure 3A). Then, a fixed-point algorithm iteratively optimises the separation vector using a cost function, with aim to maximise the sparsity of the estimated motor unit pulse train. The rationale behind this procedure is that the sum of two or more pulse trains is less sparse than a single motor unit pulse train. Therefore, the iterative optimisation of the motor unit separation vector should shift the distribution of the values of the motor unit pulse train toward a negative skew, with most of the values close to ‘0’, i.e., the noise, and a few values close to ‘1’, i.e., the discharge times (Figure 3A). This property of the motor unit pulse train perfectly matches the physiological properties of the series of motor unit discharge times, which are sparse with no more than 50 discharge times per second during submaximal isometric contractions. The fixed-point algorithm terminates once the separation vectors are sufficiently invariant between successive iterations, with respect to a pre-set upper bound, or reach 500 iterations. We implement three cost functions in MUedit: ‘logcosh’ log(cosh(x)), ‘square’ (x)^2^, and ‘skew’ (x)^3^/3. We tested the performance of the fixed-point algorithm with different initialisations of separation vectors and different cost functions with EMG signals recorded with one grid (out of four) of surface electrodes in one male and one female (Figure 3B). When initialised with the ’activity index’, the function ‘logcosh’ required more iterations than the other two functions (median: 31 iterations per motor unit, 5^th^-95^th^ percentiles: 11-500; Figure 3B). However in general, it allowed identification of a large number of motor units (32 in total for two participants at 20% of the MVC) with a highly accurate automatic estimation of the discharge times when compared to their edited version (Rate of agreement: median: 99%, 5^th^-95^th^ percentiles: 93%-100%; Figure 3B). On the contrary, the function ‘skew’ was fast to converge toward stable separation vectors (median: 16 iterations per motor unit, 5^th^-95^th^ percentiles: 11-43), but identified fewer motor units (19 in total for two participants at 20% of the MVC, all of which were already identified with logcosh, Figure 3B). The function ‘square’ represented a middle ground with fast optimisations and more motor units than the function ‘skew’ (32 in total for two participants at 20% of the MVC; Figure 3B). We also tested the impact of the initialisation on the accuracy of the decomposition using the function ‘square’, with a slightly better performance with random weights (Rate of agreement: median: 99%, 5^th^-95^th^ percentiles: 79%-100%) than with an initialisation with the ‘activity index’ (Rate of agreement: median: 98%, 5^th^-95^th^ percentiles: 86%-100%; Figure 3B). However, the user is encouraged to test the impact of these parameters on their own data and adjust them to find the optimal balance between the number of identified motor units and the speed of the decomposition according to their aims.

**Figure 3.**
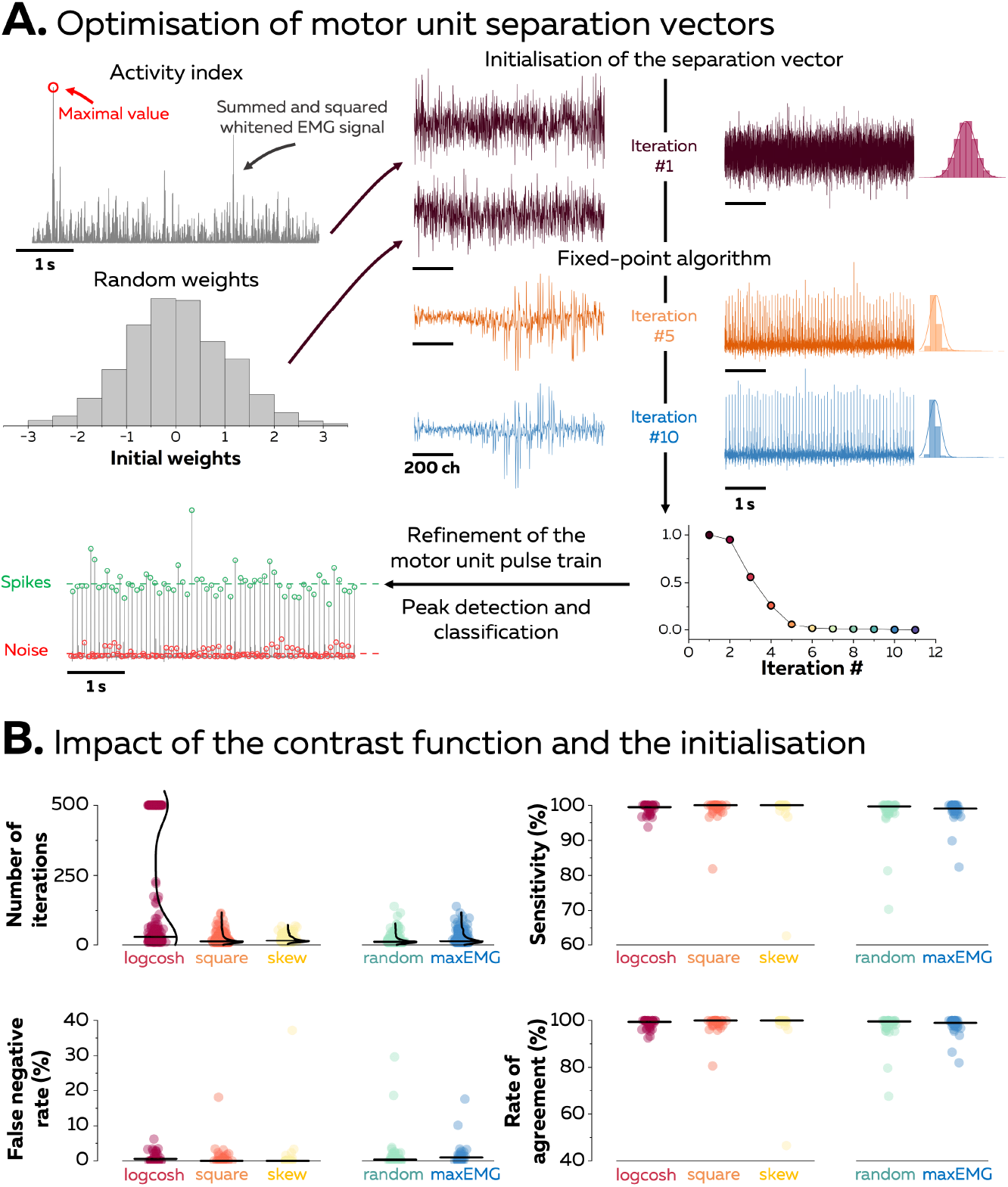
Estimation of motor unit separation vectors. (A) A fixed-point algorithm iteratively optimises separation vectors for each motor unit. The projection of EMG signals onto these separation vectors generates motor unit pulse trains, from which the series of discharge times are identified. In this example, the fixed-point algorithm terminated after 11 iterations once the separation vector varied below an imposed upper limit between successive iterations. The evolution of the separation vector, the motor unit pulse train and the distribution of its values are displayed on the right. At the end of the fixed-point algorithm, the discharge times are estimated with a peak-detection function and a K-mean classification that separate the high peaks (spikes) from the low peaks (noise). (B) We tested the impact of the initialisation of the separation vectors and the cost function used in the fixed-point algorithm on the accuracy of the automatic identification of the discharge times by comparing them to their manually edited version. Each data point is an iteration of the fixed-point algorithm (upper left panel, n = 200) or a motor unit (total n = 32, 32, and 19 for logcosh, square, and skew, respectively, n = 33 for random and max EMG), and each black line represents the median.

Once the decomposition algorithm converges to a stable separation vector and refines the motor unit pulse train, an additional step is performed to steer the algorithm towards additional sources (motor units). In the classic implementation of fastICA, this step is called ‘source deflation’ and consists of the orthogonalisation of the separation vectors to decorrelate them. Alternatively, it is possible to ‘peel-off’ the identified motor unit from the EMG signals to prevent the fixed-point algorithm to converge toward the same motor unit a second time (Figure 4A). This step consists of i) estimating the action potential waveforms for each electrode using the spike-trigger averaging method, ii) convoluting the action potentials with the series of estimated discharge times to generate a train of action potentials, and iii) subtracting this train of action potentials from the EMG signals. The next iteration is thus performed on the remaining EMG signals. We compared the performance of the decomposition with and without peeling off the identified motor units with EMG signals recorded with a grid of surface electrodes and with an intramuscular electrode array (Figure 4B). The inclusion of ‘peel-off’ increased the number of identified motor units when using the intramuscular electrode array (24 motor units in total) in comparison to ‘source deflation’ (6 motor units in total). It is worth noting that all the motor units identified with ‘source deflation’ were also identified with ‘peel-off’. The impact of ‘peel-off’ on the result of the decomposition was less significant with surface EMG signals, with only 6 additional motor units identified in comparison to the ‘source deflation’ (27 vs. 21 motor units in total). Importantly, the use of ‘peel-off’ also decreased the accuracy of the series of estimated discharge times for some motor units (Figure 4B). This is due to the misalignment of discharge times relative to the action potentials, which induces errors in the estimation of the action potential waveforms. Moreover, the presence of missed discharge times / falsely identified discharge times also induces errors when subtracting the train of action potentials from the EMG signals. Thus, the accumulation of errors across the successive iterations of the decomposition may reduce the accuracy of the last identified motor unit pulse trains, which is especially true when the accuracy of the first identified motor units is moderate. Again, the users are advised to evaluate the performance of this option on their own dataset. One way to overcome the errors of the automatic identification of series of discharge times is to additionally inspect and manually edit the results of the automatic decomposition (Del Vecchio et al., 2020), which is an option provided by MUedit.

**Figure 4.**
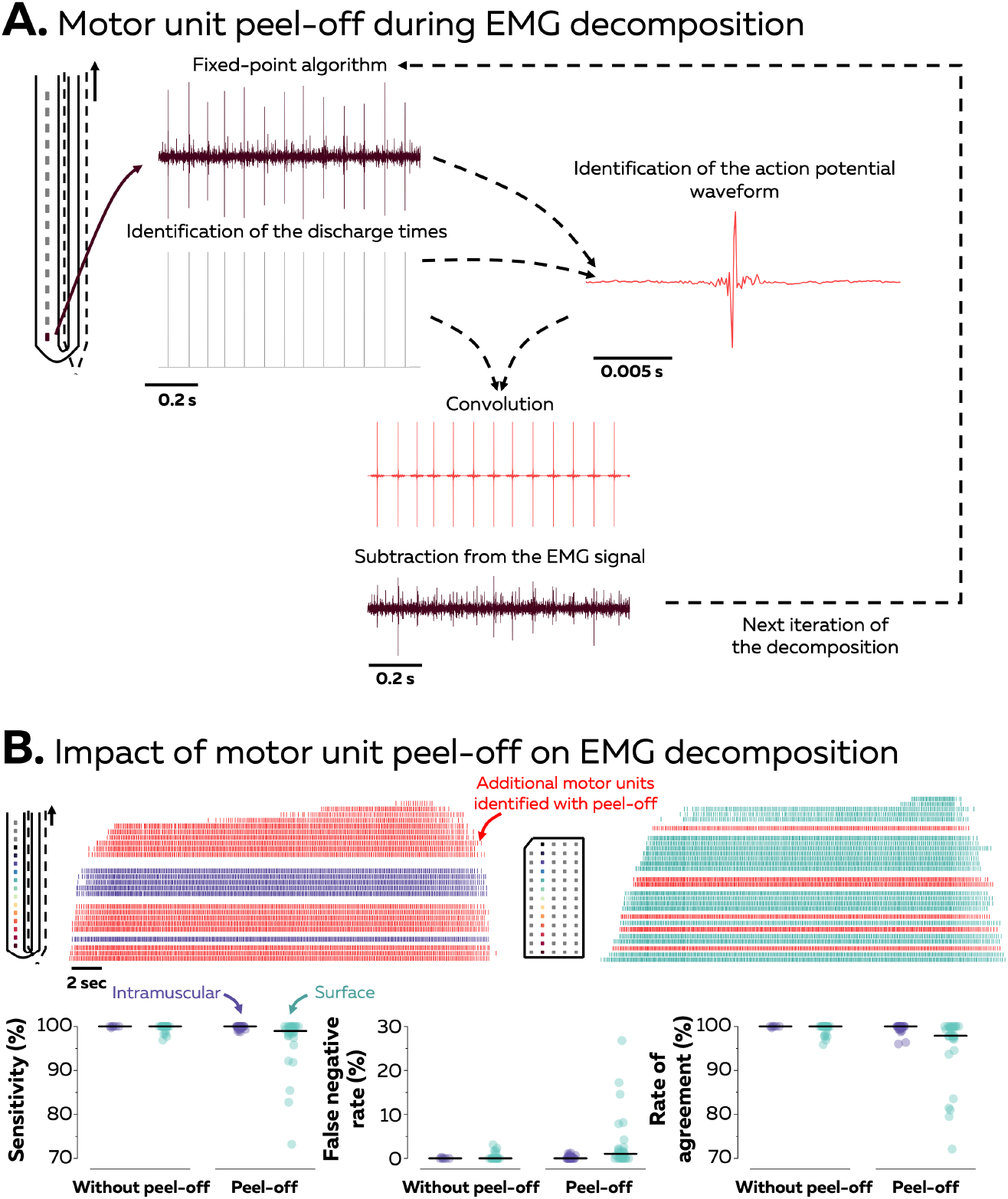
Motor unit peel-off. To prevent the decomposition algorithm from converging to the same motor unit several times, one can ‘peel-off’ the identified motor unit from the signal and run the next iteration on the remaining EMG signal. For this, the action potential waveform is identified for each electrode using the spike-trigger averaging method. As shown in Panel A, waveforms are then convoluted with the series of discharge times to generate trains of action potentials, and these trains of action potentials are subtracted from the EMG signals. Panel B depicts raster plots of motor units identified from intramuscular and surface EMG signals. Each tick is a discharge time. The series of discharge times exclusively identified when ‘peel-off’ is active are displayed in red. We measured the accuracy of the automatic identification of the discharge times by comparing them to their manually edited version. Each data point is a motor unit, and each black line represents the median.

## Manual editing of motor unit pulse trains

The results provided by the decomposition algorithm need to be assessed for accuracy (Holobar et al., 2014). The silhouette value measures the normalised distance between the spikes and the noise and can be used to assess the reliability of the motor unit pulse trains (Negro et al., 2016). Previous studies recommended to only consider motor units which exhibit a silhouette value higher than 0.9. However, it is important to keep in mind that even when the silhouette value reaches this threshold, the motor unit pulse train must be visually inspected to check for errors of the automatic identification of discharge times. To speed up manual editing, we implemented an option to automatically perform the first steps of the manual editing (Figure 5). Specifically, i) outliers can be automatically removed from the series of discharge times, ii) separation vectors can be recalculated and reapplied over the entire EMG signals, and then iii) outliers can be automatically removed a second time. This is a simplified and more practical version than the method recently proposed by Clarke and Farina (2021). The performance of this automatic step is displayed on Figure 5. Generally, the automatic recalculation of separation vectors works well when motor units are associated with a sufficiently high silhouette value. After the automatic update of the motor unit pulse train, the user can quickly check the motor unit pulse train and the series of discharge times with minimal manual editing. On the contrary, the automatic recalculation of separation vectors does not improve the accuracy of the motor unit pulse trains with numerous initial errors in the estimation of discharge times. For these motor units, visual inspection and manual editing can effectively compensate for inaccuracies of the automatic decomposition.

**Figure 5.**
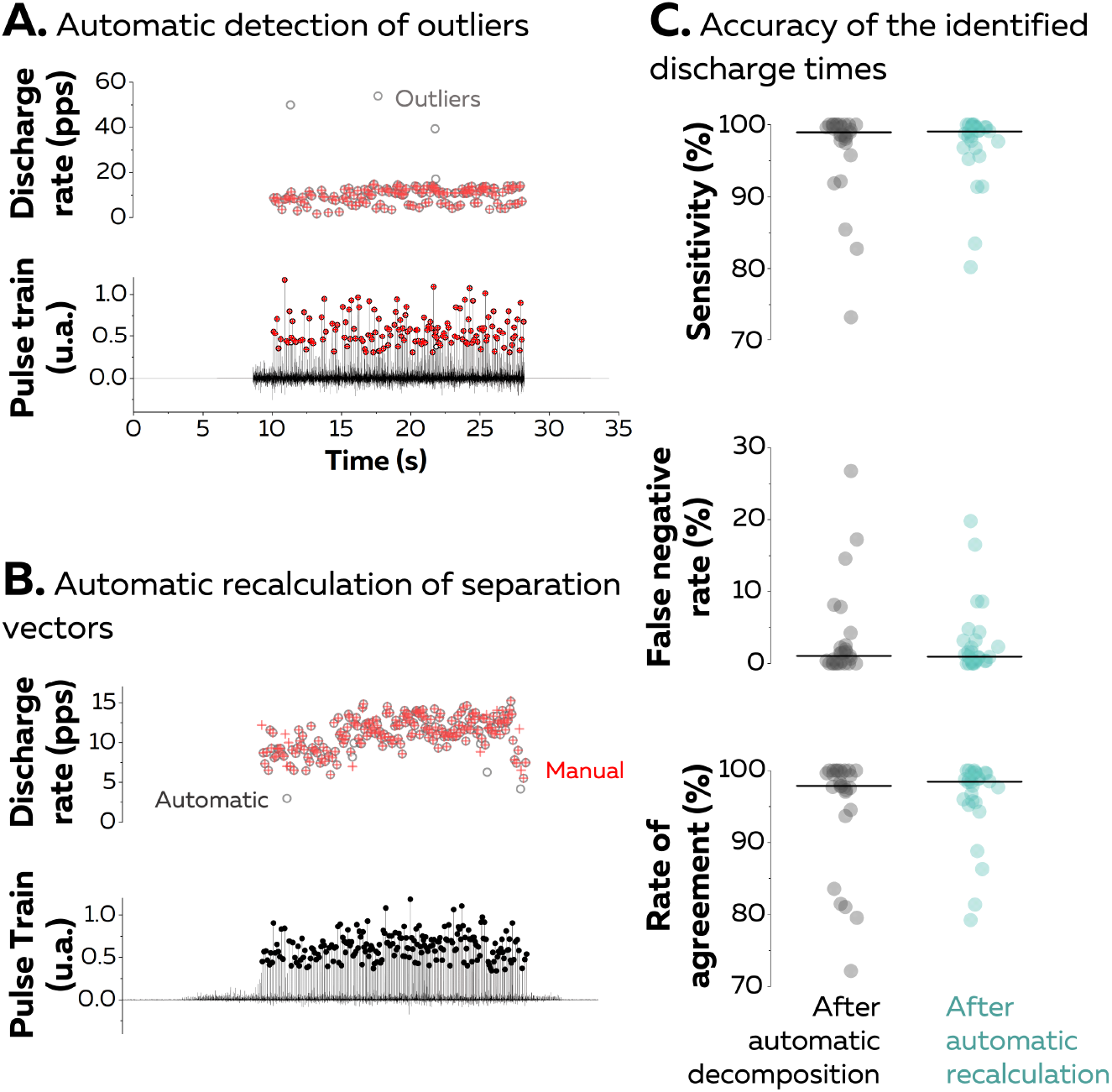
Automatic update of motor unit pulse trains. (A) The first step consists of removing the discharge times causing erroneous discharge rates, i.e., higher than the average discharge rate + three standard deviations. On this graph, one can also observe drops in discharge rates due to missed discharge times. (B) The recalculation of the motor unit pulse trains and the automatic identification of the discharge times is similar to the procedure used in automatic decomposition. Specifically, the discharge times are estimated with a peak-detection function and a K-mean classification that separate the high peaks (spikes) from the low peaks (noise). The removal of outliers is then repeated a second time. On the right, the plots display the accuracy of the automatic identification of the refined discharge times by comparing them to their manually edited version. Each data point is a motor unit, and each black line represents the median.

The manual editing is supported by MUedit and consists of i) removing the spikes causing erroneous discharge rates (outliers), ii) adding the discharge times clearly separated from the noise, iii) recalculating the separation vector, iv) reapplying the separation vector on the entire EMG signals, and v) repeating this procedure until the selection of all the discharge times (Figure 6). The manual editing of potential missed discharge times and falsely identified discharge times should not be immediately accepted. Instead, the procedure is consistently followed by the application of the updated motor unit separation vector on the entire EMG signals to generate a new motor unit pulse train. The manual edition should only be accepted when the silhouette value increases following this operation or stay well above the threshold of 0.9. A more extensive description of the manual editing of motor unit pulse trains can be found in Del Vecchio et al. (2020) and Hug et al. (2021).

**Figure 6.**
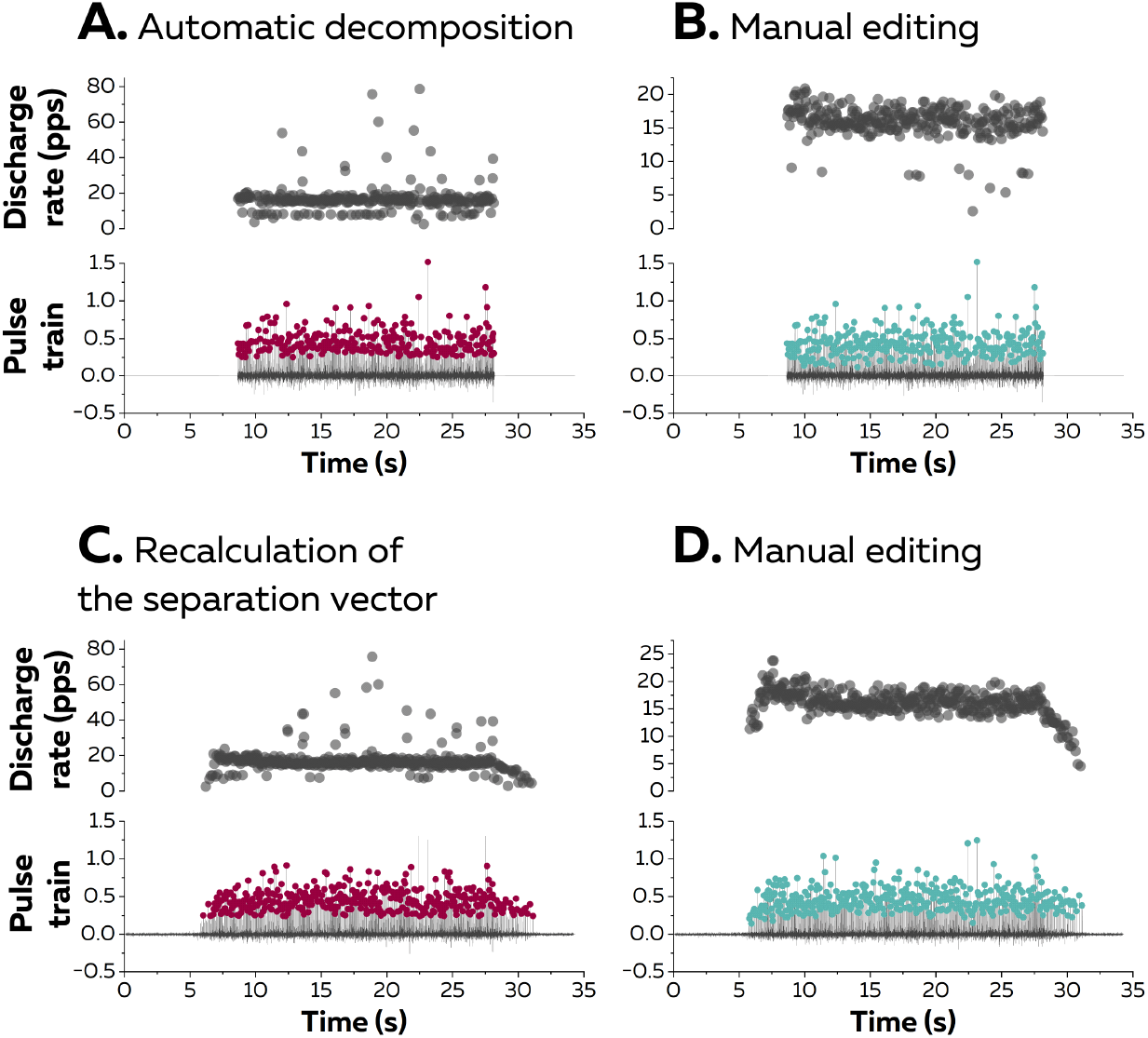
Manual editing of motor unit pulse trains. (A) After the automatic decomposition, falsely identified discharge times cause erroneous discharge rates while missed discharge times cause drops in discharge rates. (B) One can manually remove or add these discharge times if they are clearly identifiable. (C) After this manual editing, the separation vector is recalculated, and the motor unit pulse train is updated. (D) The automatic detection of discharge times is manually corrected another time, and this iterative process stops once the series of discharge times is completed.

Once all the series of discharge times are visually inspected and manually edited, the last step consists of removing the duplicates in the rare case where they were not automatically removed after the automatic decomposition. For this purpose, the estimated series of discharge times from pairs of motor units are aligned using a cross-correlation function to account for a potential delay due to the propagation time of action potentials along the fibres. Then, two discharge times are considered as common when they occur within a time interval of 0.5 ms, and motor units are considered as duplicates when they have at least 30% of their identified discharge times in common (Holobar et al., 2010).

We recently observed that using several grids of surface electrodes over the same muscle can dramatically increase the number of identified motor units (Caillet et al., 2023). For example, the separate decomposition of EMG signals recorded from four grids of 64 electrodes positioned over the Tibialis Anterior enabled us to identify 80 instead of 22 motor units in a male participant and 42 instead of 10 motor units in a female participant (Figure 7). In this case, the user should also remove the duplicates between grids, as the surface potentials arising from the same motor unit can be detected by multiple grids of electrodes.

**Figure 7.**
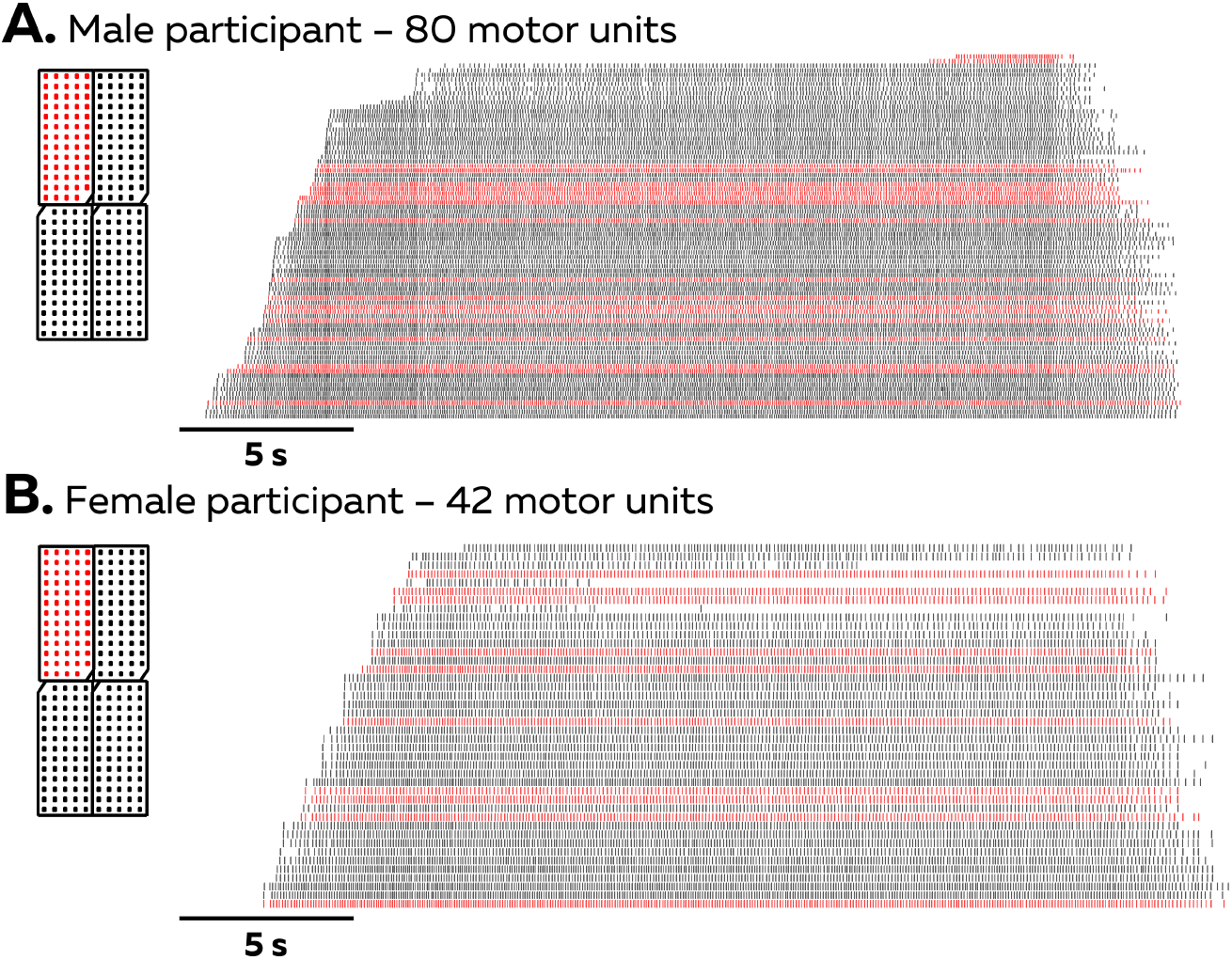
Decomposition of multiple grids. We recorded and decomposed surface EMG signals from four grids of 64 electrodes positioned next to each other with an interelectrode distance of 4 mm. After removing the duplicates, we identified 80 (A) and 42 (B) motor units for the male and the female participant, respectively. The raster plots display the discharge times for all the motor units. The red ticks are the discharge times of the motor units identified from the grid #1, while the grey ticks are the discharge times of the motor units identified from all the other grids.

## Limitations of EMG decomposition

The decomposition algorithm currently implemented in MUedit can only estimate the series of discharge times during stationary conditions, where the activation and the geometry of the muscle are stable. In practice, this means that we progressively lose the discharge times when the action potential waveforms change over time. One way to overcome this limitation is to iteratively recalculate the separation vectors on successive windows (Chen et al., 2020, Glaser and Holobar, 2019, Oliveira and Negro, 2021). This process could be partly automatised by optimising the separation vectors according to an additional variable related to these changes, such as the joint angle (Glaser and Holobar, 2019), and performing a rigorous visual inspection and manual editing of the motor unit pulse trains.

Moreover, the algorithm quickly fails when the force level is very high, which is due to the additional recruitment of motor units that overlap with the identified motor units. This leads to merged pulse trains where the discharge times of the different motor units are indistinguishable. Manual editing can sometime solve this issue when the height of the peaks clearly differs between motor unit pulse trains.

Another limitation of the decomposition algorithm is the potential misalignment of some estimated discharge times with the occurrence of the motor unit action potential. This is highlighted in this tutorial paper by the rate of agreement of 94% between the series of discharge times of a motor unit simultaneously detected from surface and intramuscular EMG signals, though we identified the same number of discharge times in both decompositions (Figure 2). One way to overcome these small misalignments is to realign the discharge times with the onset of each motor unit action potential identified by spike-trigger segmenting the single-differential EMG signal (Ibáñez et al., 2021). Again, this process should be associated with a rigorous visual inspection of the action potential waveforms and the detected onsets.

## Conclusion

We introduced the open-source software MUedit accompanied by a documented code, a user manual, and dataset with aim to standardise the use of the current state-of-the-art of EMG decomposition. The algorithm was first validated on simulated surface and intramuscular EMG to prove the high accuracy of the estimation of series of motor unit discharge times. We then presented how the initialisation of the separation vector, the contrast function used in the fixed-point algorithm, and the deflation approach impact the result of the decomposition. Finally, we emphasised the importance of visual inspection and manual editing to guarantee the accuracy of the decomposition. We hope that the development of this tool will expand the use of EMG decomposition in experimental and clinical settings and encourage users to contribute to the code on the online repository.

## Supporting information

User manual

## Supplemental data availability

The entire data set (raw and processed data), codes, and a user manual of the software are available at https://github.com/simonavrillon/MUedit for a version coded with Matlab (version 2022b, The MathWorks, Inc, USA). A version of the software coded with Python (Python 3.9.15, Python Software Foundation, USA) is available at https://github.com/ciaragibbs/MUEdit_Python.

## Notes

**Conflict of interests:** The authors declare no competing financial interests.

**Funding resources.** Dario Farina is supported by the European Research Council Synergy Grant NaturalBionicS (contract #810346), the EPSRC Transformative Healthcare, NISNEM Technology (EP/T020970), and the BBSRC, “Neural Commands for Fast Movements in the Primate Motor System” (NU-003743). François Hug is supported by a fellowship from the French government, through the UCAJEDI Investments in the Future and by the National Research Agency (ANR) with the reference number ANR-15-IDEX-01.

https://github.com/simonavrillon/MUedit

### Competing Interest Statement

The authors have declared no competing interest.

## References

Adrian ED, Bronk DW.The discharge of impulses in motor nerve fibres: Part II. The frequency of discharge in reflex and voluntary contractions. J Physiol. 1929;67:i3–151.

Armstrong JB, Rose PK, Vanner S, Bakker GJ, Richmond FJ.Compartmentalization of motor units in the cat neck muscle, biventer cervicis. J Neurophysiol. 1988;60:30–45.

Basmajian JV, Stecko G. A new bipolar electrode for electromyography. J Appl Physiol (1985). 1962;17:849–.

Caillet AH, Avrillon S, Kundu A, Yu T, Phillips ATM, Modenese L, et al. Larger and denser: an optimal design for surface grids of EMG electrodes to identify greater and more representative samples of motor units. bioRxiv. 2023:2023.02.18.529050.

Chen C, Ma S, Sheng X, Farina D, Zhu X. Adaptive Real-Time Identification of Motor Unit Discharges From Non-Stationary High-Density Surface Electromyographic Signals. IEEE Trans Biomed Eng. 2020;67:3501–9.

Chen M, Zhou P. A Novel Framework Based on FastICA for High Density Surface EMG Decomposition. IEEE Trans Neural Syst Rehabil Eng. 2016;24:117–27.

Chung B, Zia M, Thomas K, Michaels JA, Jacob A, Pack A, et al. Myomatrix arrays for high-definition muscle recording. bioRxiv. 2023:2023.02.21.529200.

Clarke AK, Farina D. Deep Metric Learning with Locality Sensitive Angular Loss for Self-Correcting Source Separation of Neural Spiking Signals. arXiv preprint arXiv:211007046.2021.

de Luca CJ.The use of surface electromyography in biomechanics. J Appl Biomech. 1997;13:135–63.

Del Vecchio A, Holobar A, Falla D, Felici F, Enoka RM, Farina D. Tutorial: Analysis of motor unit discharge characteristics from high-density surface EMG signals. J Electromyogr Kinesiol. 2020;53:102426.

Duchateau J, Enoka RM. Human motor unit recordings: origins and insight into the integrated motor system. Brain Res. 2011;1409:42–61.

Farina D, Holobar A. Characterization of Human Motor Units From Surface EMG Decomposition. Proceedings of the Ieee. 2016;104:353–73.

Farina D, Merletti R, Enoka RM. The extraction of neural strategies from the surface EMG. J Appl Physiol (1985). 2004;96:1486–95.

Farina D, Negro F, Gazzoni M, Enoka RM. Detecting the unique representation of motor-unit action potentials in the surface electromyogram. J Neurophysiol. 2008a;100:1223–33.

Farina D, Negro F, Muceli S, Enoka RM. Principles of Motor Unit Physiology Evolve With Advances in Technology. Physiology (Bethesda). 2016;31:83–94.

Farina D, Yoshida K, Stieglitz T, Koch KP. Multichannel thin-film electrode for intramuscular electromyographic recordings. J Appl Physiol (1985). 2008b;104:821–7.

Formento E, Botros P, Carmena JM. Skilled independent control of individual motor units via a non-invasive neuromuscular-machine interface. J Neural Eng. 2021;18.

Gazzoni M, Farina D, Merletti R. A new method for the extraction and classification of single motor unit action potentials from surface EMG signals. J Neurosci Methods. 2004;136:165–77.

Glaser V, Holobar A. Motor Unit Identification From High-Density Surface Electromyograms in Repeated Dynamic Muscle Contractions. IEEE Trans Neural Syst Rehabil Eng. 2019;27:66–75.

Holobar A, Farina D. Blind source identification from the multichannel surface electromyogram. Physiol Meas. 2014;35:R143–65.

Holobar A, Minetto MA, Botter A, Negro F, Farina D. Experimental analysis of accuracy in the identification of motor unit spike trains from high-density surface EMG. IEEE Trans Neural Syst Rehabil Eng. 2010;18:221–9.

Holobar A, Minetto MA, Farina D. Accurate identification of motor unit discharge patterns from high-density surface EMG and validation with a novel signal-based performance metric. J Neural Eng. 2014;11:016008.

Holobar A, Zazula D. Multichannel Blind Source Separation Using Convolution Kernel Compensation. IEEE Transactions on Signal Processing. 2007;55:4487–96.

Hug F, Avrillon S, Del Vecchio A, Casolo A, Ibanez J, Nuccio S, et al. Analysis of motor unit spike trains estimated from high-density surface electromyography is highly reliable across operators. J Electromyogr Kinesiol. 2021;58:102548.

Ibáñez J, Del Vecchio A, Rothwell JC, Baker SN, Farina D. Only the fastest corticospinal fibers contribute to beta corticomuscular coherence. J Neurosci. 2021.

Jiang X, Liu X, Fan J, Ye X, Dai C, Clancy EA, et al. Open Access Dataset, Toolbox and Benchmark Processing Results of High-Density Surface Electromyogram Recordings. IEEE Trans Neural Syst Rehabil Eng. 2021;29:1035–46.

Konstantin A, Yu T, Le Carpentier E, Aoustin Y, Farina D. Simulation of Motor Unit Action Potential Recordings From Intramuscular Multichannel Scanning Electrodes.IEEE Trans Biomed Eng. 2020;67:2005–14.

LeFever RS, De Luca CJ. A procedure for decomposing the myoelectric signal into its constituent action potentials--Part I: Technique, theory, and implementation. IEEE Trans Biomed Eng. 1982;29:149–57.

Liddell EGT, Sherrington CS. Recruitment and some other features of reflex inhibition. Proc Biol Sci. 1925;97:488–518.

McGill KC, Lateva ZC, Marateb HR. EMGLAB: an interactive EMG decomposition program. J Neurosci Methods. 2005;149:121–33.

Muceli S, Poppendieck W, Holobar A, Gandevia S, Liebetanz D, Farina D. Blind identification of the spinal cord output in humans with high-density electrode arrays implanted in muscles. Science advances. 2022;8:eabo5040.

Muceli S, Poppendieck W, Negro F, Yoshida K, Hoffmann KP, Butler JE, et al. Accurate and representative decoding of the neural drive to muscles in humans with multi-channel intramuscular thin-film electrodes. J Physiol. 2015;593:3789–804.

Nawab SH, Wotiz RP, De Luca CJ. Decomposition of indwelling EMG signals. J Appl Physiol (1985). 2008;105:700–10.

Negro F, Muceli S, Castronovo AM, Holobar A, Farina D. Multi-channel intramuscular and surface EMG decomposition by convolutive blind source separation. J Neural Eng. 2016;13:026027.

Oliveira AS, Negro F. Neural control of matched motor units during muscle shortening and lengthening at increasing velocities. J Appl Physiol (1985). 2021;130:1798–813.

Rau G, Disselhorst-Klug C, Silny J. Noninvasive approach to motor unit characterization: muscle structure, membrane dynamics and neuronal control. J Biomech. 1997;30:441–6.

