## Supplementary material for "Tutorial on MUedit: An open-source software for identifying and analysing the discharge timing of motor units from electromyographic signals": User manual

### MUedit: a step-by-step user manual

#### Introduction:

MUedit is a Matlab app that decomposes electromyographic (EMG) signals recorded from multiple electrodes into individual motor unit pulse trains using fast independent component analysis (fastICA). You can easily adjust the parameters of the algorithm to the specificity of your experimental settings. After the decomposition, you can visualise and edit the output of the fastICA, i.e., the motor unit pulse trains. We provide here a step-by-step protocol to facilitate the implementation of MUedit in any experimental settings.

#### Before starting MUedit:

##### Software requirements.

MUedit works on any modern computer running Matlab. The current version of MUedit has been developed on Matlab R2022b and tested on a laptop equipped with an Apple M1 Max chip and 64 GB of RAM. However, we successfully ran MUedit on multiple Windows, Linux, and MacOs computers with versions of Matlab ranging from 2018a to 2023a.

MUedit requires three Matlab toolboxes: signal processing, image processing, and statistics and machine learning. You need to install them before running the app for the first time.

##### MUedit installation.

Up-to-date versions of MUedit are uploaded on GitHub in the following repository https://github.com/simonavrillon/MUedit . The repository is structured with one folder 'lib' containing all the matlab functions needed to run MUedit, and four matlab scripts containing either:

1. the full code (MUedit_exported.m),
2. the code + the design of the app (MUedit.mlapp),
3. an installation file to have the app directly runnable from the Matlab's app library (MUedit.mlappinstall),
4. the full code of the decomposition algorithm (HDEMGdecomposition.m).

*Option #1: MUedit_exported.m*

To run MUedit_exported.m, you first need to add the full folder with the library of functions and the main code to Matlab's path. i) Go to the 'Home' tab, 'Environment' table, and click on 'Set Path'; ii) Click on 'Add with Subfolders…', find the folder with all the scripts, and click on 'Open'; iii) Click on 'Save', go to the 'Editor' tab, click on 'Open', find the script MUedit_exported.m, and click on 'Run'.

*Option #2: MUedit.mlapp*

To run MUedit.mlapp, you need to open the app editor. i) Go to the 'Apps' tab and click on 'Design App'; ii) Click on 'Open', find the script MUedit.mlapp and open it; iii) Go to the 'Designer' tab and click on 'Run'.

*Option #3: MUedit.mlappinstall*

Before the first use, you need to install the MUedit app. Open the script MUedit.mlappinstall and click on 'Install' in the pop-up window. For all the following uses, open Matlab, go to the 'Apps' tab, find the app in the apps library and click on the shortcut to run the app.

*Option #4: HDEMGdecomposition.m*

To run HDEMGdecomposition.m, you first need to add the full folder with the library of functions and the main code to Matlab's path. i) Go to the 'Home' tab, 'Environment' table, and click on 'Set Path'; ii) Click on 'Add with Subfolders…', find the folder with all the scripts, and click on 'Open'; iii) Click on 'Save', go to the 'Editor' tab, click on 'Open', find the script HDEMGdecomposition.m, and click on 'Run'.

#### Description of the user interface:

MUedit has two main panels: 'Decomposition' and 'Edition'.

On the 'Decomposition' panel, you can adjust the options and parameters of the decomposition algorithm to optimise its performance. Once the parameters are ready, you can hit the 'Start' button to start the decomposition.

On the 'Edition' panel, you can import the decomposed files, visualise the motor unit pulse trains, and manually edit and correct the discharge times. You can plot the results and save the edited version of your motor unit pulse trains.

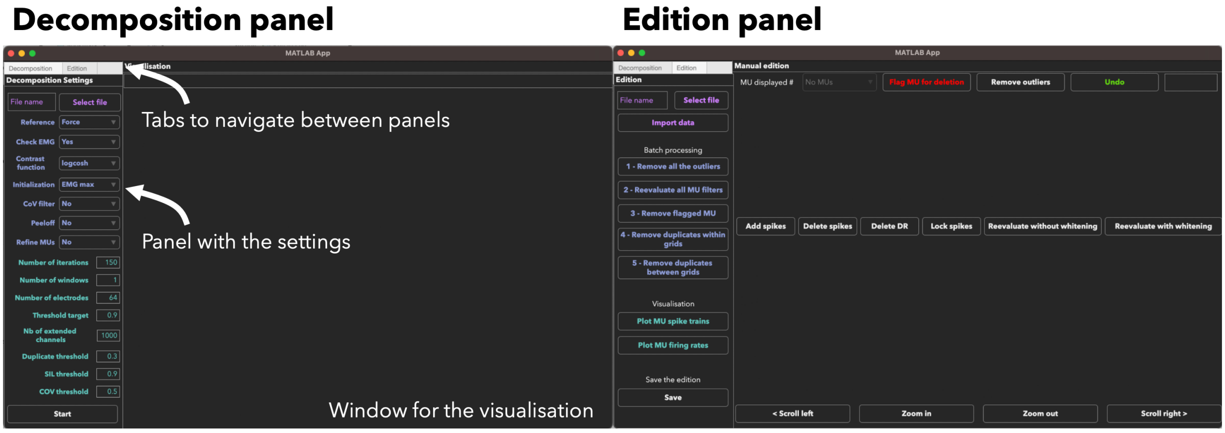


**Figure 1.** Overview of the panels from the MUedit app.

#### Overview of the approach:

An EMG signal represents the sum of trains of action potentials from all the active motor units within the recorded muscle volume. During stationary conditions, e.g., isometric contractions, the train of motor unit action potentials can be modelled as the convolution of series of discrete delta functions, representing the discharge times, and motor unit action potentials (Figure 2A). The EMG decomposition with blind source separation consists of inversing the generative model of electromyographic signals by estimating the series of discharge times for each active motor unit (Figure 2B). When EMG signals are recorded with an array of electrodes, the shape of the recorded potential of each motor unit differs across electrodes. This is due to 1) the varying conduction velocity of action potentials among the muscle fibres, and 2) the location/depth of the muscle fibres that belong to each motor unit relatively to the electrodes, which impact the low pass filtering effect of the tissue on the recorded potential. Therefore, the EMG signals recorded with an array of electrodes can be considered as an instantaneous mixture of the original motor unit spike trains and their delayed versions.

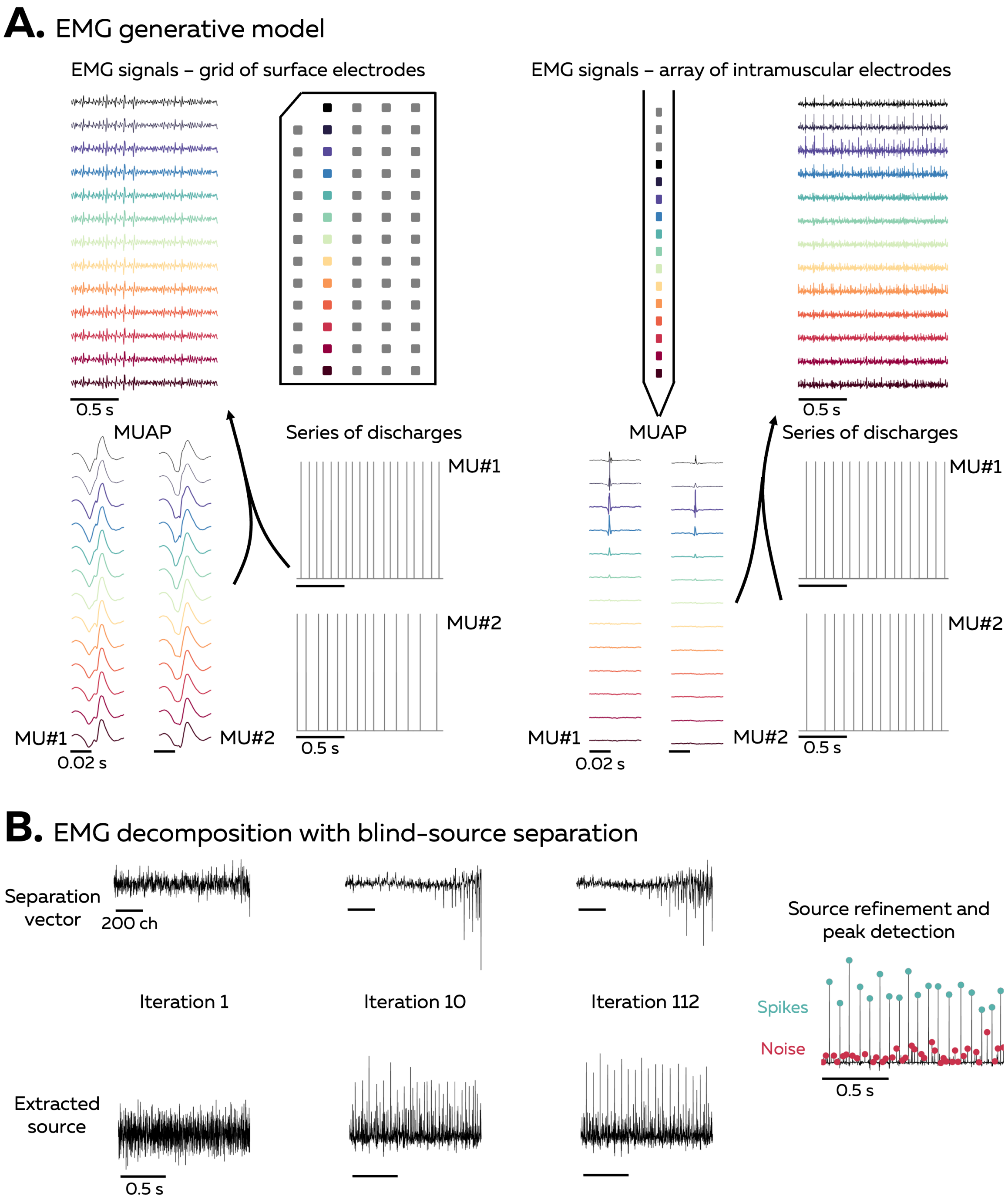


**Figure 2.** Overview of the approach.

Increasing the number and spatial distribution of recording electrodes increases the likelihood that each motor unit will have a unique motor unit action potential profile (shape), i.e., a temporal and spatial profile that differs from all the other active motor unit within the recorded volume. The uniqueness of motor unit action potential profiles is necessary for the blind source separation to accurately estimate the motor unit discharge times. Our software uses a fast independent component analysis (fastICA) to optimise the separation vector for each motor unit (Figure 2B). The projection of EMG signals onto the separation vectors will generate motor unit pulse trains, from which we estimate the discharge times of each motor unit (Figure 2B).

**<u>Overview of the decomposition algorithm and the associated functions:</u>**

1) The monopolar EMG signals collected during the experimental session are imported in the software (openOTBplus.m), displayed on the screen such that you can remove channels with artifacts or bad signal-to-noise ratio (if the option is selected), and reformatted (formatsignalHDEMG.m).
2) The signals are filtered with i) an adaptive notch filter that remove the frequencies with abnormal peaks (notchsignals.m) and ii) a band pass filter (20-500 Hz for surface EMG and 100-4400 Hz for intramuscular EMG) (bandpassingals.m).
3) The EMG signals are extended by adding delayed versions of each channel (extend.m).
4) The EMG signals are then demeaned (demean.m) and spatially whitened (pcaesig.m; whiteesig.m) to make them uncorrelated and of equal power.
5) A fixed-point algorithm is applied to identify the motor unit pulse trains (fixedpointalg.m). In this algorithm, a contrast function is iteratively applied to estimate a separation vector that maximised the level of sparseness of the motor unit pulse train.
6) At this stage, the motor unit pulse train contained high peaks (i.e., the spikes from the identified motor unit) and low peaks (i.e. spikes from other motor units and noise). High peaks are separated from low peaks and noise using peak detection and K-mean classification with two classes: 'spikes' and 'noise' (getspikes.m). The peaks from the class with the highest centroid are considered as spikes of the identified motor unit.
7) A second algorithm refines the estimation of the discharge times by iteratively recalculating the separation vector and repeating the steps with peak detection and K-mean classification until the coefficient of variation of the inter-spike intervals is minimised (minimizeCOVISI.m).
8) The accuracy of each estimated pulse train is assessed by computing the silhouette (SIL) value between the two classes of spikes identified with K-mean classification (calcSIL.m). When the SIL exceeds a predetermined threshold, the motor unit is saved, and optionally peeled-off (peeloff.m) from the signal to run the next iteration on the residual of the EMG signals.

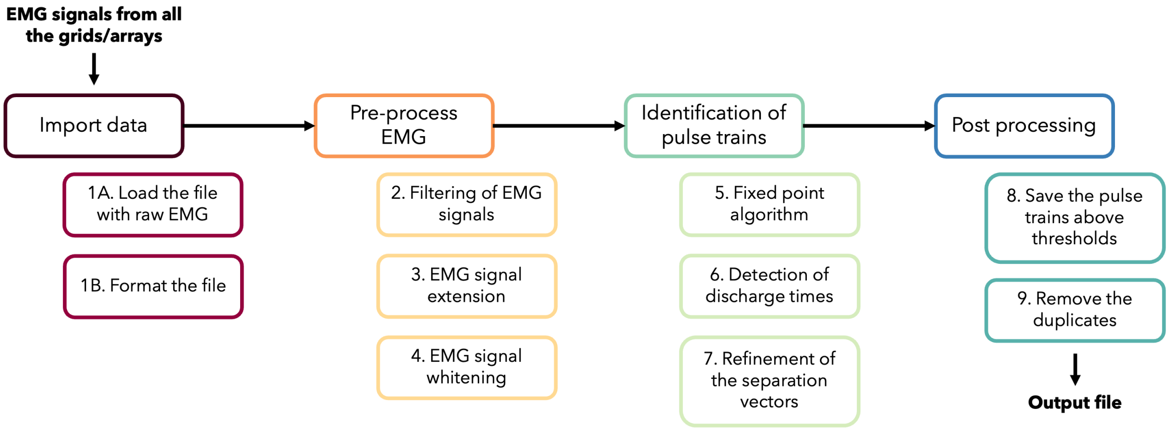


**Figure 3.** Main steps of the decomposition process.

**Format of EMG signals to import:**

You can import two types of files: i) '.otb+' files recorded with the software OT BioLab+ and ii) '.mat' files formatted for MUedit.

When recording from OT BioLab+, go to Acquisition, Setup, and configure electrodes type (e.g., 'GR08MM1305') and the muscles (e.g., 'Tibialis Anterior'). The sampling frequency should be set at a minimum of 2,048 Hz. MUedit will automatically import the grid and muscle names, the force, and the target together with the EMG signals from the '.otb+' file. However, it is worth noting that the automatic import only works when all the signals are recording with grids. If you also record signals from the 16 inputs channels, you should follow the procedure for importing '.mat. files.

When importing a '.mat' file, create a structure called signal (Figure 4) and add the following variables:

- **signal.data** is a matrix of a size *nb channels* × *time* where EMG signals from all the grids are vertically concatenated. For example, EMG signals recorded from two grids of 64 electrodes over 10 seconds at a sampling frequency of 2,048 Hz would generate a matrix signal.data of a size 128 × 20480.
- **signal.fsamp** is the sampling frequency.
- **signal.nChan** is the number of recorded channels.
- **signal.ngrid** is the number of grids/arrays used to record EMG signals.
- **signal.gridname** is an array of cells containing the names of the grids as strings. For example, EMG signals recorded with 2 grids of 64 electrodes, 13 × 5, interelectrode distance 8mm from OT Bioelettronica would generate an array of two cells, with 'GR08MM1305' in each cell. If you used different grids or your own system, you must create a name of grids here, and add the configuration of the grid in the function formatsignalHDEMG.m (see below)
- **signal.muscle** is an array of cells containing the names of the muscles as strings. For example, EMG signals recorded with 2 grids from the tibialis anterior and the soleus would generate an array of two cells, with 'Tibialis Anterior' and 'Soleus' in each cell.

Optional:

- **signal.target** is a vector of a size *1* × *time* containing the force/torque target displayed to the participant.
- **signal.path** is a vector of a size *1* × *time* containing the force/torque produced by the participant.

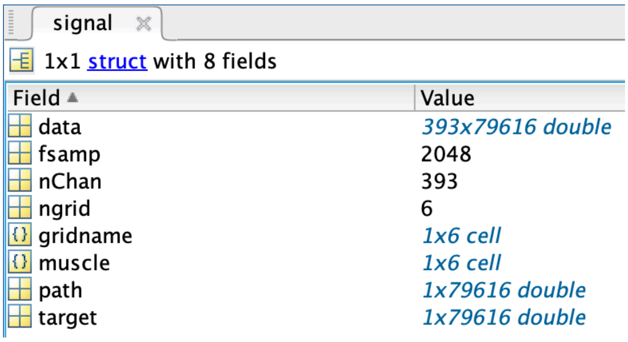


**Figure 4.** Matlab structure "*signal*" to import data in MUedit.

If you use another system, or a custom-made grid/array of electrodes, you must also add its configuration in the function formatsignalHDEMG.m. For this, you will need to add in the function `switch gridname` the case with the name of your grid (Figure 5) and the following variables:

- **ElChannelMap** is a matrix or a vector with the position of each channel within the grid.
- **discardChannelsVec** is a vector of zeros of a size *1 × number of electrodes in the grid*.
- **IED** is the interelectrode distance, e.g., 4 for 4mm.
- **emgtype** is the type of signals. 1 being for EMG signals recorded with surface electrodes, 2 for EMG signals recorded with intramuscular electrodes.

```
case 'GR04MM1305'
    ElChannelMap = ([1 25 26 51 52; ...
                    1 24 27 50 53; ...
                    2 23 28 49 54; ...
                    3 22 29 48 55; ...
                    4 21 30 47 56; ...
                    5 20 31 46 57; ...
                    6 19 32 45 58 ; ...
                    7 18 33 44 59; ...
                    8 17 34 43 60; ...
                    9 16 35 42 61; ...
                    10 15 36 41 62; ...
                    11 14 37 40 63; ...
                    12 13 38 39 64]);
    discardChannelsVec = zeros(64,1);
    IED = 4;
    emgtype = 1;
```

**Figure 5.** Code example for the grid 'GR04MM1305' in the function formatsignalHDEMG.m.

Of note, the current version of MUedit includes the following grid names sold by OTBioelettronica: 'GR04MM1305', 'ELSCH064NM2', 'GR08MM1305', 'GR10MM0808'.

After recording the EMG signals from OT Biolab+ or creating the '.mat' file with the structure signal, go to MUedit, click on Select file, and open either the '.otb+' or the '.mat' file'.

#### Parameters of the decomposition:

Before starting the decomposition, you can adjust a few options and parameters to optimise the performance of the algorithm.

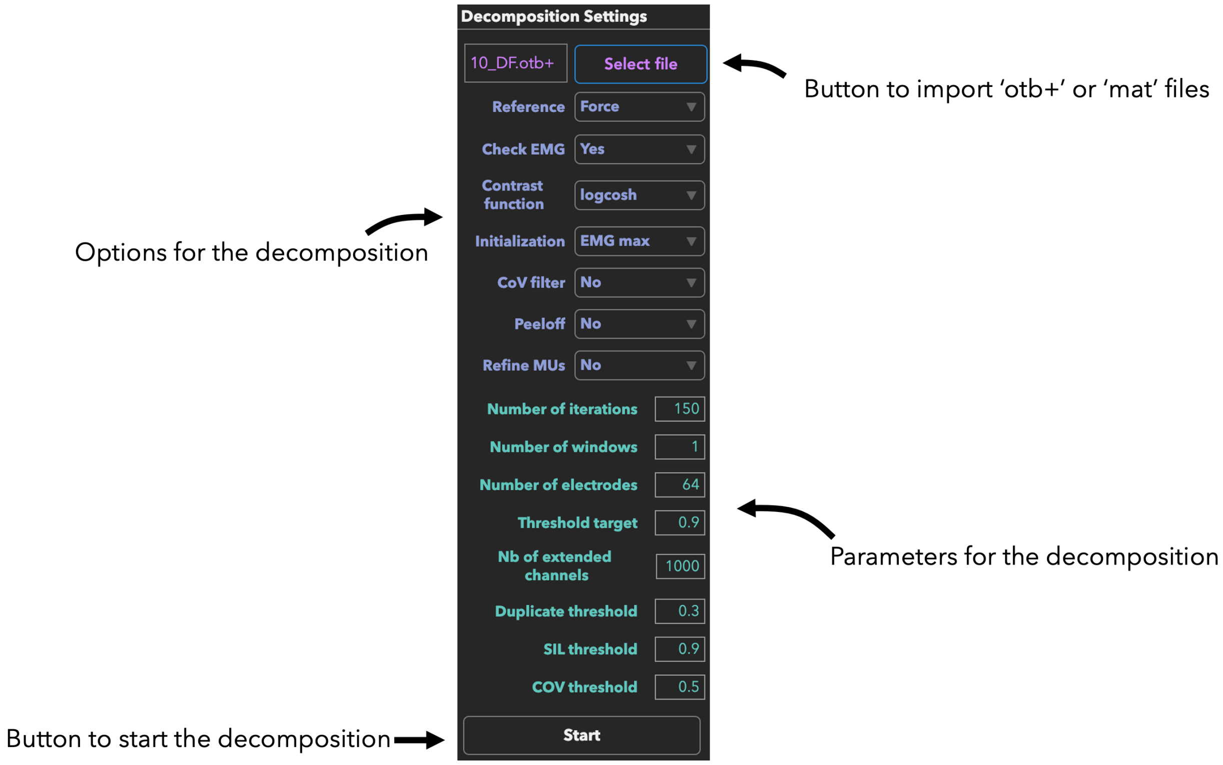


**Figure 6.** Panel of EMG decomposition.

- **Reference**: if you used the trapezoidal feedback from OT Biolab+ or imported your target and force signals in the structure *signal* as *signal.target* and *signal.path* (see section 'Format of EMG signals to import'), you can select 'Force'. The algorithm will automatically segment the EMG signals to decompose the section(s) for which the actual force is above a threshold that you can freely chose ("Threshold target" in the parameters for decomposition). For example, if you have a plateau set at 10% of the maximal force, and you want to automatically decompose the EMG signals above 9% of the maximal force, set the value of Threshold target to 0.9 as 90% of the level of the target. Alternatively, if you want to manually select the sections of the signals to decompose, select 'EMG amplitude'. At the beginning of the decomposition, you will be asked to select a region interest over a plot displaying the rectified EMG signal of half the channels and its average value.
- **Check EMG**: if you want to visually check the EMG signals and manually remove channels with low signal-to-noise ratio and/or artifacts, select 'Yes'. If you want to automatically decompose all the channels, select 'No'.
- **Contrast function**: you can select the contrast function used in the fixed-point algorithm that optimise separation vectors for each motor unit. You have three choices: 'logcosh', 'square', and 'skew'.
- **Initialisation**: you can choose how to initialise the separation vectors in the fixed-point algorithm. If you select 'EMG max', the separation vectors will be initialised as the EMG values at the time instance where the sum of the squared EMG values over all the channels is maximal. If you select 'Random', the separation vectors will be initialised with random values.

- **CoV filter**: If you want to only keep motor unit pulse trains with stable and constant discharge times, you can select 'Yes' and add a threshold to keep motor units which exhibit a coefficient of variation of their inter-spike intervals below the fixed COV threshold. If not, select 'No'.
- **Peeloff**: If you want to remove the trains of action potentials of reliable identified motor units from the EMG signals and run the following iteration on the residual EMG, Select 'Yes'. If not, select 'No'. Reliable identified motor units have pulse trains above the silhouette threshold.
- **Refine MUs**: you can choose to automatically "clean" the identified motor unit pulse trains. If you select 'Yes', the algorithm iteratively deletes discharge times such that discharge rates above a threshold (mean + 3 standard deviations) are removed. This loop stops once the coefficient of variation of the inter-spike intervals is below 0.3. After this, the separation vector is updated based on the new discharge times and applied over the entire signal. The algorithm finally iteratively deletes discharge times to remove discharge rates above the threshold for a second time.
- **Number of iterations**: Number of iterations performed by the decomposition algorithm per grid and window (see below).
- **Number of windows**: Number of windows for which the signal is decomposed. If you selected 'Force' as reference, the number of windows corresponds to the number of segments above the threshold. If you selected 'EMG amplitude as reference, this is the number of regions of interest that you want to select/analyse.
- **Number of electrodes**: Number of electrodes per grid/array.
- **Threshold target**: Threshold to automatically segment the target of force displayed to the participant. This value is only used if you selected 'Force' as reference.
- **Nb of extended channels**: Number of channels after signal extension.
- **Duplicate threshold**: Percentage of common discharge times for two motor units to be considered as duplicates. 0.3 means 30% of the total number of discharge times from the motor unit with the highest number of discharge times.
- **SIL threshold**: Only motor unit pulses trains with a silhouette value above this threshold will be retained.
- **COV threshold**: If you selected 'Yes' to COV filter, only motor unit pulses trains with a coefficient of variation of their inter-spike intervals below this threshold will be retained.

**EMG decomposition:**

After clicking on 'Start', a few manual steps may be completed to visually check EMG signals or select regions of interests for the decomposition (see above).

If you selected 'Yes' to the option "Check EMG", columns of EMG signals will successively appear on the screen (Figure 7). Enter the number of the rows to be removed (if multiple rows, use space in between) and hit 'Ok'. If you want to keep all the rows, hit 'Ok'. This step ends once all the columns from all the grids have been checked.

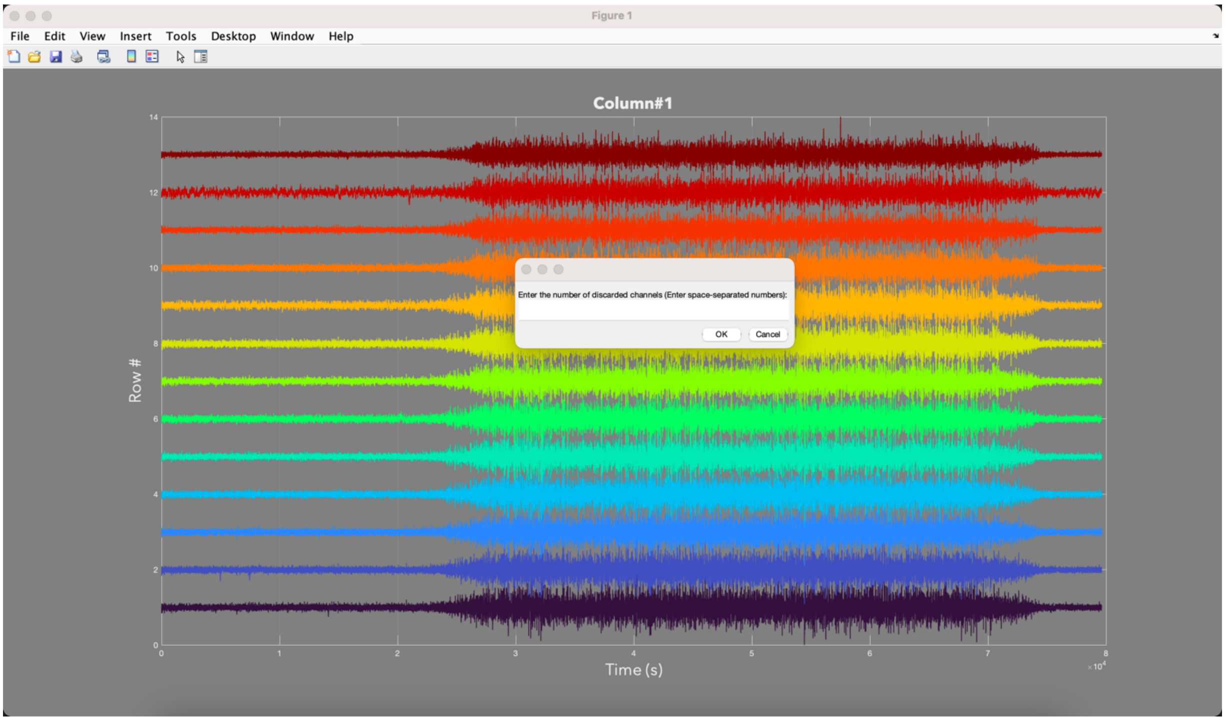


**Figure 7.** Window to visually check the EMG signals.

If you selected 'EMG amplitude' as reference, you must select as many regions of interest as the number of windows set in the EMG decomposition panel. For this, drag and drop the rectangle box over the segment of EMG amplitude displayed on the upper plot (Figure 8).

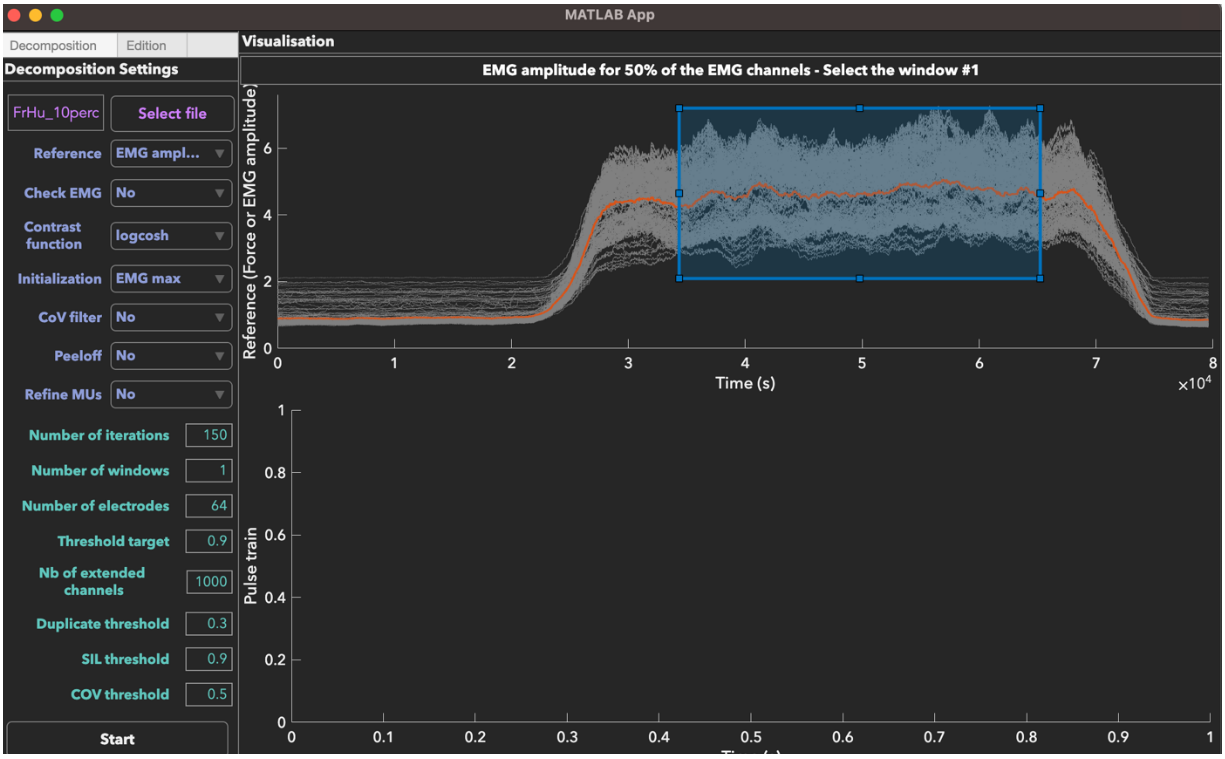


**Figure 8.** Window to select the region of interest over the EMG amplitude.

After these steps, the EMG decomposition starts. The app displays on the upper plot the segment selected with the reference. It also displays the pulse train and the identified discharge times on the bottom plots. Information about the number of the grid, the number of the iteration, the silhouette value, and the coefficient of variation of the inter-spike intervals is displayed on the top of the app and is refreshing at each iteration (Figure 9).

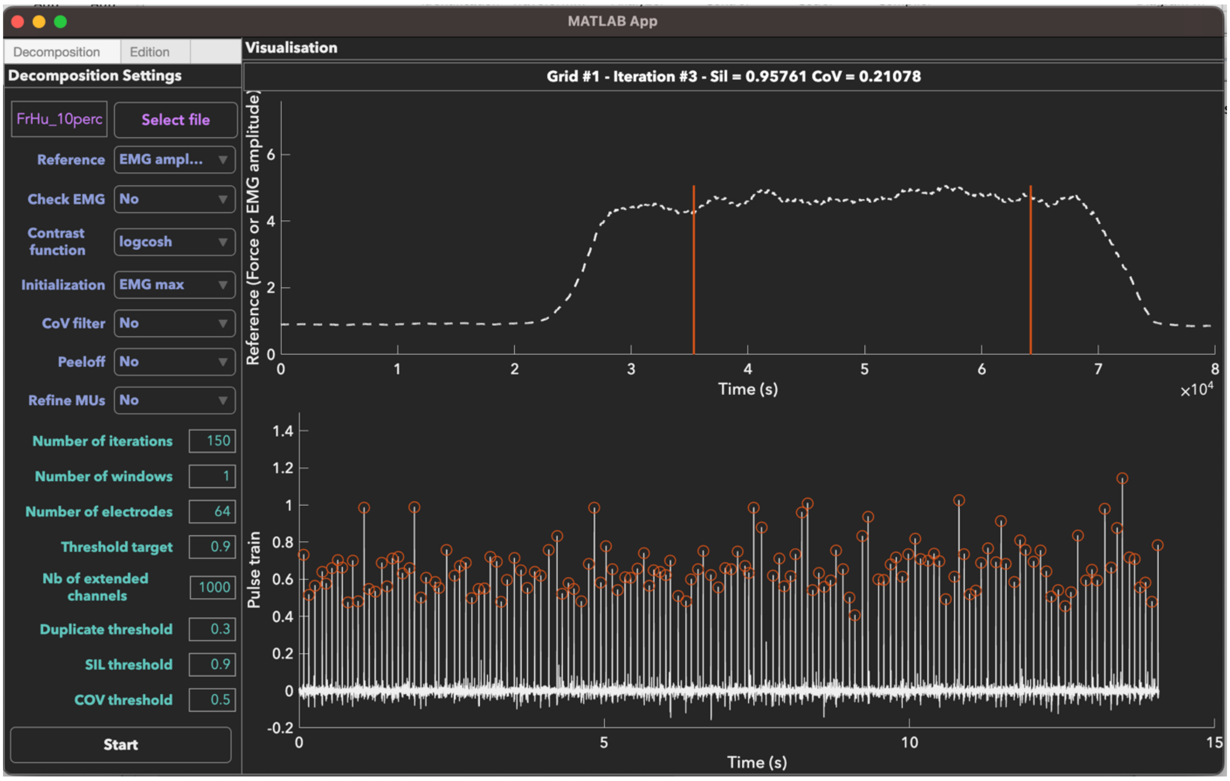


**Figure 9.** Window during the decomposition. On the top plot, the dashed line represents the reference (here the average EMG amplitude). The red lines display the limits of the region of interest. On the bottom plot, the white trace represents the motor unit pulse train, and the red scatters the identified discharge times for the current iteration.

At the end of the decomposition, a '.mat' file is automatically saved with a structure "parameters" which contains all the parameters of the decomposition and a structure "signal" which contains all the data. In addition of the initial variables gathered in "signal", two variables with the output of the decomposition are added:

- **signal.Pulsetrain** is an array of cells containing the motor unit pulse trains for each grid/array of electrodes. The size of each cell is *nb motor units* × *time*.
- **signal.Dischargetimes** is a set of cells of a size *nb grids* × *nb motor units* containing the discharge times of each identified motor unit.

#### Manual edition of motor unit pulse trains:

It is recommended to visually check and manually edit the motor unit pulse trains and the identified discharge times. The manual edition aims to correct errors from the peak separation between the two classes (spikes and noise) using K-mean classification. Specifically, the following steps are performed:

- identifying and removing the spikes of lower quality.
- updating the motor unit separation vector and re-applying it over a portion of the signal.
- adding the new spikes recognized as motor unit discharge times.

It is worth noting that the manual identification of potential missed spikes and false positives is never arbitrarily accepted. Indeed, it is always followed by the update and the re-application of the separation vector of the motor unit, which reveals whether the manual edition should be accepted or rejected based on the change in silhouette value. For additional information, you can follow the guidelines reported in Hug et al. (2021) and Del Vecchio et al., (2020).

Hug F, Avrillon S, Del Vecchio A, Casolo A, Ibanez J, Nuccio S, Rossato J, Holobar A & Farina D. (2021). Analysis of motor unit spike trains estimated from high-density surface electromyography is highly reliable across operators. J Electromyogr Kinesiol 58, 102548.

Del Vecchio A, Holobar A, Falla D, Felici F, Enoka RM & Farina D. (2020). Tutorial: Analysis of motor unit discharge characteristics from high-density surface EMG signals. J Electromyogr Kinesiol 53, 102426.

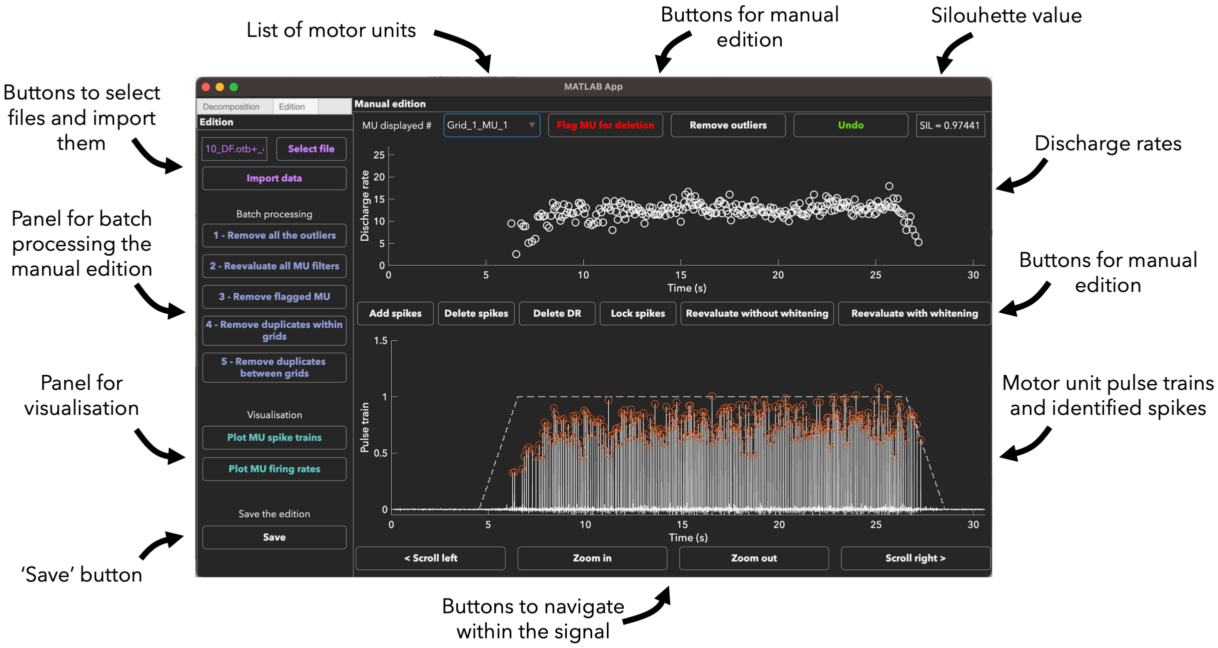


**Figure 9.** Description of the interface of MUedit for manual edition.

To start with manual edition, click on Select file, select the '.mat' file generated by the decomposition algorithm, Open, and Click on Import Data. To navigate between grids and motor units, you can go to the drop-down menu on the upper left.

Below are the functions of the buttons dedicated to manual edition:

- **Flag MU for deletion**: if the motor unit pulse train is not reliable and need to be deleted, click on this button. It won't delete the MU immediately, but it will be deleted once you click on 'remove flagged MU' (left panel).
- **Remove outliers**: identify the instantaneous discharge rates above a threshold (mean discharge rate + 3 standard deviation) and remove the discharge times causing this discharge rate with the lowest height.
- **Undo**: cancel the last action of manual edition.
- **Add spikes**: drag and drop a region of interest around the spikes you want to select. You can select multiple spikes at the same time.
- **Delete spikes**: drag and drop a region of interest around the spikes you want to remove. You can remove multiple spikes at the same time.
- **Delete DR**: drag and drop a region of interest around the instantaneous discharge rates you want to remove. It will remove the discharge time causing this discharge rate with the lowest height. You can remove multiple instantaneous discharge rates at the same time.
- **Re-evaluate without whitening**: update the motor unit separation vector based on the identified discharge times within the window displayed on the screen and reapply it over the extended EMG signals of the same window. It is worth noting that the identification of the discharge times will be updated using K-mean classification.
- **Re-evaluate with whitening**: update the motor unit separation vector based on the identified discharge times within the window displayed on the screen and reapply it over the extended and whitened EMG signals of the same window. It is worth noting that the identification of the discharge times will be updated using K-mean classification.
- **Lock spikes**: If you want to re-evaluate the separation vectors without an automatic selection of the discharge times, click on this button. It will lock the identified discharge times such that they won't be removed in the re-evaluation process. You will need to click on this button every time you want to lock the discharge times and re-evaluate. Of note, using this option leads to a more subjective manual edition.

Typically, manual edition starts by identifying missed spikes or potential false positives by looking at the variation in discharge rate on the upper plot. On Figure 10-1, you can observe a few drops in discharge rates due to the presence missed spikes. You can use the button 'Add spikes' to drag and drop a region of interest around the missed spikes (Figure 10-2) and repeat this for all the spikes (Figure 10-3). After this, you can click on Re-evaluate without whitening (fast) or with whitening (slower) to update the motor unit separation vector and apply it over the EMG signals within the window. The colour of the scatters will depend on the changes in silhouette value. **Red** means that the silhouette value changes by more than -4, **orange** by between -4 and -2, **blue** by between 0 and +2, and **green** above 2. Generally, manual edition should improve the silhouette value (green scatters).

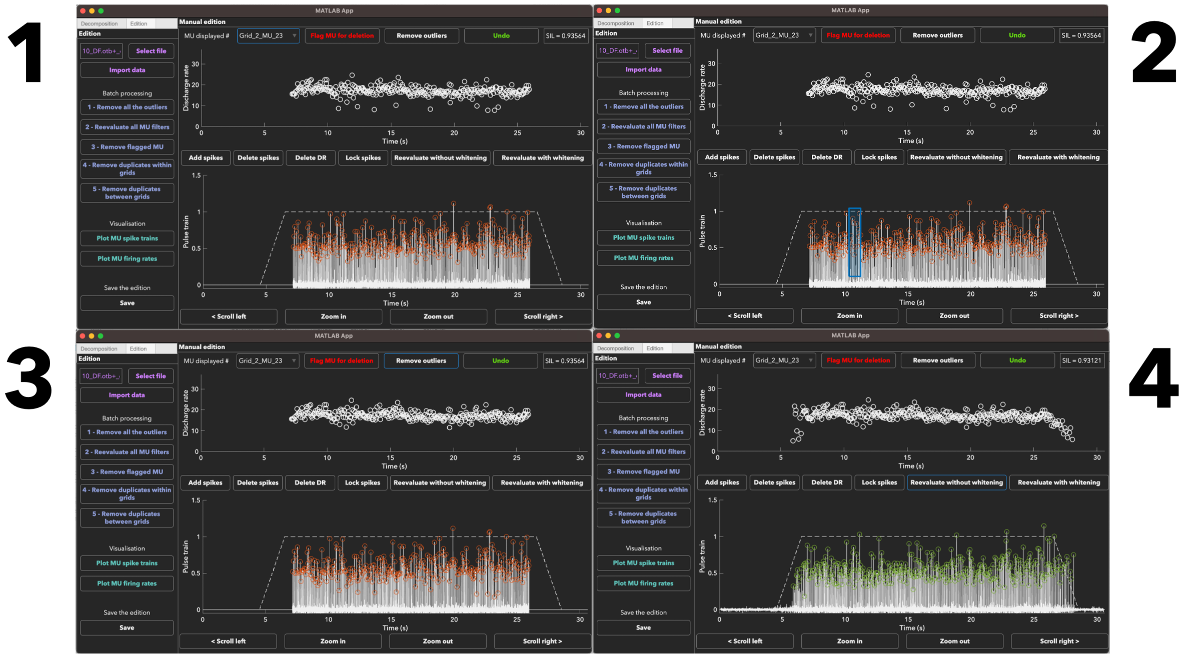


**Figure 10.** Successive steps of manual edition. 1) identification of the missed spikes (discharge rates drop). 2) Click on 'Add spikes' and drag and drop a region of interest around the spikes to selected. 3) Motor unit pulse train and discharge rates with all the spikes selected. 4) Re-evaluate the motor unit separation vector and apply it over the entire EMG signals. The green scatters mean that the silhouette value improved with manual edition.

To speed up manual edition, you can batch process some processing steps, such as removing the discharge rates considered as outliers, or updating all the motor unit separation vectors and reapplying them all at once. For this, click on 'Remove all the outliers' or 'Reevaluate all MU filters' on the left panel and this will be applied on all motor units.

At the end of the edition, it is recommended to check for duplicates between motor units within the grids (if you used one grid per muscle) and between grids (if you used multiple grids on the same muscles). We consider two or more motor units as duplicates when they have at a significant percentage of their identified discharge times in common. This percentage is set as 'Duplicate threshold' in the panel of the decomposition tab, with 0.3 being 30% of discharge times in common. The motor unit pulse trains are first aligned using a cross-correlation function to account for a potential delay due to the conduction time along the fibres. Then, two discharge times are considered as common when they occurred within a time window of 0.5 ms. When a duplicate is found, the motor unit kept in the analysis is the one with the lowest coefficient of variation of inter-spike intervals. The buttons for removing the duplicates are in the left panel.

You can visualise the all the motor units with two types of plots (Figure 11). The first one displays the raster plots of each grid with ticks for each discharge time. The second one displays the motor unit discharge rates smoothed with a Hanning window of one second.

Raster plots of discharge times Smoothed discharge rates

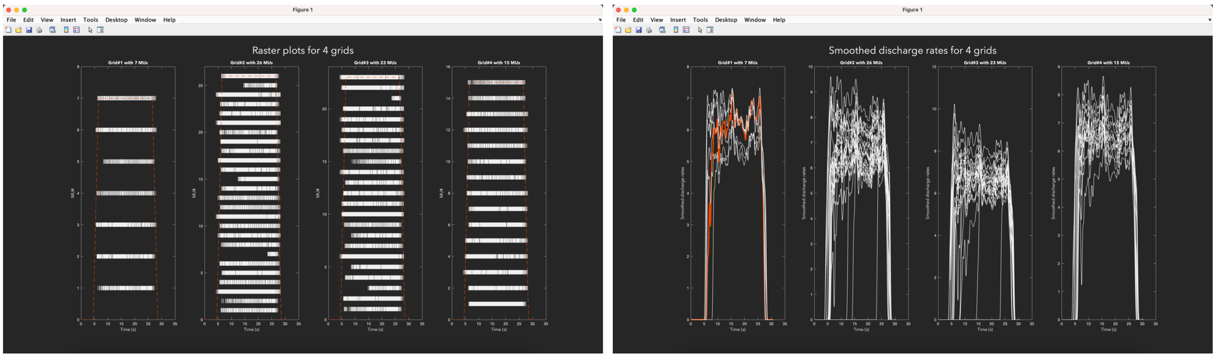


**Figure 11.** Visualisation of all the motor units in MUedit

At the end of the session, click on 'Save' to save a '.mat' file with the edited motor unit pulse trains and discharge times. It will keep the same file name with the addition of '_edited' at the end. In the file, a new structure named edition will appear with the same variables as in signal, but with the edited pulse trains (edition.Pulsetrain) and discharge times (edition.Dischargetimes). To further process the same file, you can load this edited file. You will automatically resume the edition where you stopped it.

#### Citation and technical support.

For technical assistance and support, please contact:

Dr. Simon Avrillon
Sir Michael Uren Hub
Imperial College London
86 Wood Ln
London W12 0BZ
ORCID: 0000-0002-2226-3528
